# A folded conformation of MukBEF and Cohesin

**DOI:** 10.1101/464701

**Authors:** Frank Bürmann, Byung-Gil Lee, Thane Than, Ludwig Sinn, Francis J O’Reilly, Stanislau Yatskevich, Juri Rappsilber, Bin Hu, Kim Nasmyth, Jan Löwe

## Abstract

Structural maintenance of chromosomes (SMC)-kleisin complexes organize chromosomal DNAs in all domains of life, where they have key roles in chromosome segregation, DNA repair and regulation of gene expression. They function through topological entrapment and active translocation of DNA, but the underlying conformational changes are largely unclear. Using structural biology, mass spectrometry and cross-linking, we investigated the architecture of two evolutionarily distant SMC-kleisin complexes: proteobacterial MukBEF and eukaryotic cohesin. We show that both contain a dynamic coiled-coil discontinuity, the elbow, near the middle of their arms that permits a folded conformation. Bending at the elbow brings into proximity the hinge dimerization domain and the head/kleisin module, situated at opposite ends of the arms. Our findings favor SMC activity models that include a large conformational change in the arms, such as a relative movement between DNA contact sites during DNA loading and translocation.

## Introduction

All organisms organize and maintain enormous chromosomal DNA molecules whose contour lengths exceed cellular dimensions by orders of magnitude. Hence both regulation of gene expression and accurate chromosome segregation require a high degree of spatial organization. Structural maintenance of chromosomes (SMC)-kleisin complexes are ancient machines that help to organize chromosome superstructure in bacteria, archaea and eukaryotes (Hirano, 2016). They are essential for chromosome segregation in many bacteria and are retained in the set of 438 protein-coding genes from an organism with an artificially reduced genome (Britton et al., 1998; Hutchison et al., 2016; Jensen and Shapiro, 1999; Niki et al., 1991). Similarly, SMC-kleisin complexes are essential for both mitosis and meiosis in eukaryotes (Guacci et al., 1997; Hirano and Mitchison, 1994; Klein et al., 1999; Michaelis et al., 1997; Pebernard et al., 2004; Saka et al., 1994).

At the core of SMC-kleisin complexes is a conserved tripartite protein ring, composed of an SMC homo- or hetero-dimer, bridged by a kleisin subunit (Bürmann et al., 2013; Haering et al., 2002; Onn et al., 2007; Woo et al., 2009; Zawadzka et al., 2018). SMC proteins are elongated molecules containing an ABC-type ATPase head and a hinge dimerization domain at opposite ends of an approximately 50 nm long intramolecular and antiparallel coiled-coil arm (Anderson et al., 2002; Diebold-Durand et al., 2017; Haering et al., 2002; Melby et al., 1998; Niki et al., 1992). The kleisin asymmetrically connects the two heads of an SMC dimer and contains binding sites for additional KITE or HAWK subunits (Palecek and Gruber, 2015; Wells et al., 2017).

A widely conserved and possibly fundamental aspect of SMC-kleisin activity is the ability to entrap chromosomal DNA within their ring structure (Cuylen et al., 2011; Gligoris et al., 2014; Wilhelm et al., 2015). DNA entrapment is the molecular basis for sister chromatid cohesion by the cohesin complex, and might also be used by cohesin, condensin and bacterial Smc-ScpAB to organize DNA into large loops. Loading of DNA into the complex is thought to involve transient opening of a ring interface in cohesin (Gruber et al., 2006; Murayama and Uhlmann, 2015), and is likely mediated by the SMC arms in Smc-ScpAB (Bürmann et al., 2017).

The second possibly universal aspect of SMC-kleisin activity is their translocation along DNA. Cohesin and bacterial SMC-kleisin complexes associate with chromosomes in a manner that requires ATP binding and they redistribute or translocate from initial loading sites to adjacent regions dependent on ATP hydrolysis (Badrinarayanan et al., 2012; Hu et al., 2011; Minnen et al., 2016; Wang et al., 2018). Translocation of bacterial Smc-ScpAB coincides with progressive linking of distant chromosomal loci *in vivo*, indicating that the complex might actively extrude DNA loops (Wang et al., 2017; Wang et al., 2018). Recently, ATPase dependent DNA translocation and loop extrusion reactions have been reconstituted using purified condensin *in vitro* (Ganji et al., 2018; Terakawa et al., 2017). These findings support the notion that SMC-kleisin complexes are motor proteins that use the ATPase activity of their SMC subunits to track along DNA, and some, by doing so, might actively organize chromosomes by building up loops (Alipour and Marko, 2012; Marsden and Laemmli, 1979; Nasmyth, 2001).

To explore how the core activities of SMC-kleisin complexes might be implemented on a structural level, we have investigated the architecture of two representative complexes that are separated by a billion years of evolution: MukBEF from *E. coli* and cohesin from budding yeast. We find that both complexes contain a bendable structural coiled-coil discontinuity in their arms that allows them to interconvert between extended and folded conformations, in the latter bringing the hinge dimerization domain close to the head/kleisin module. Our findings show that SMC proteins have the capacity for a large conformational transformation, and provide the basis for investigating long-distance domain movements during DNA loading and translocation reactions.

## Results

### A folded conformation of MukBEF and cohesin

MukBEF is a diverged SMC-kleisin complex that serves as an essential chromosome organization machine in *E. coli* (Badrinarayanan et al., 2012; Lioy et al., 2018; Woo et al., 2009; Yamanaka et al., 1996; Yamazoe et al., 1999). The complex comprises the SMC protein MukB, the kleisin MukF and the KITE protein MukE. We co-overexpressed the MukBEF subunits in *E. coli* and prepared the complex by a multi-step procedure that yielded purified material without extra residues on any of the subunits (Fig. 1a). The purified complex eluted as a single peak in size exclusion chromatography (SEC) (Fig. 1b) and was analyzed by negative stain electron microscopy (EM) immediately after elution from the column (Fig. 1c). Although subject to heterogeneity, most particles had a characteristic double cherry-like shape, composed of a two-lobed density (the MukB head/MukEF module) from which a stalk emerged (the MukB arms). Surprisingly, many particles possessed a stalk length of about 24 nm, roughly half of what is expected for an extended MukB arm consisting of canonical coiled-coil segments. As evident from partially extended particles, this conformation was caused by folding at a kink close to the center of the MukB arms. We propose to refer to this kink as the “elbow”, as it connects the upper and lower parts of the arms (Fig. 1c). Fully extended particles were also observed, but were less apparent. Using reference-free 2D image classification we obtained class averages for the conformationally less heterogeneous closed form (Fig. 1c). Class averages displayed the MukB head/MukEF module as a bow-tie shaped density with a central bridge and showed a clear signal for the folded arm with the elbow at its vertex. We also observed the presence of the elbow by cryo-electron microscopy imaging (cryo-EM) of a distantly related (~26 % sequence identity) MukBEF complex embedded in vitreous ice, without the use of particle support or contrast agent **(Supplementary Fig. 1**).

**Figure 1.**
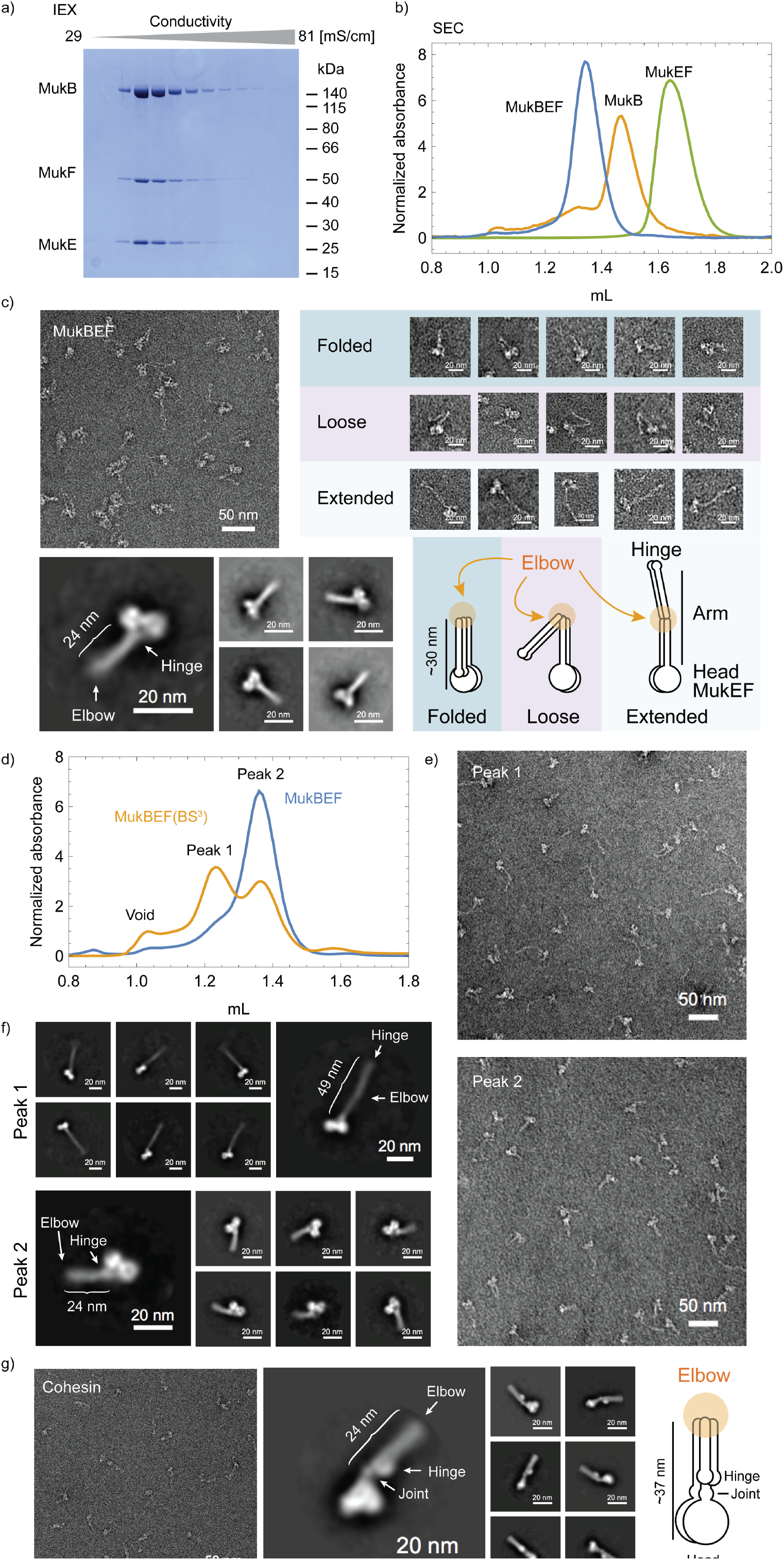
Folded conformation of MukBEF and cohesin. **(a)** Purification of MukBEF. Elution of the MukBEF complex from a Q ion exchange (IEX) column. Peak fractions were separated by SDS-PAGE and stained with Coomassie Blue. **(b)** Size-exclusion chromatography (SEC) of the MukBEF complex, MukB and MukEF. Proteins were separated on Superose 6 Increase. **(c)** Negative stain EM of native MukBEF. A typical field of view is shown in the top left panel. Particle instances for observed conformations are shown in the top right panel. 2D class averages for the folded conformation are shown on the bottom left, using a circular mask of 640 Å. **(d)** Cross-linking of MukBEF with BS3. SEC profiles for native and cross-linked material are shown. **(e)** Negative stain EM microscopy of BS3 cross-linked MukBEF. Typical fields of view for particles from SEC peak 1 and SEC peak 2 are shown. **(f)** Negative stain 2D class averages for extended (top) and folded (bottom) conformations, using circular masks of 948 Å and 640 Å, respectively. Data was collected from samples of peak 1 and peak 2 of the SEC shown in (d). **(g)** Negative stain EM of BS3 cross-linked cohesin. A typical field of view is shown on the left. Class averages using a circular mask of 500 Å are shown in the middle panel.

We noticed the presence of a considerable fraction of what appeared to be broken particles on the negative stain EM grids, possibly caused by the grid preparation procedure. To decrease heterogeneity, we subjected the *E. coli* MukBEF to mild cross-linking with the amine-reactive compound BS3 (bis(sulfosuccinimidyl)suberate). This treatment caused the complex to elute from SEC in two major peaks: one at a retention volume similar to native material, and one eluting earlier, indicative of an increased hydrodynamic radius (Fig. 1d). Electron micrographs of material eluting close to native material revealed particles mostly in the folded conformation with significantly reduced heterogeneity (Fig. 1e). The faster eluting fraction migrated differently from reconstituted MukBEF doublets (Petrushenko et al., 2006) (Supplementary Fig. 2a) and was, interestingly, enriched for singlet particles in an extended conformation (Fig. 1e). We readily obtained 2D class averages for both open and closed conformations of BS3 cross-linked MukBEF using this fractionation approach **(Fig. 1f**). In the averages, the MukB elbow is positioned at a clearly resolved near-central position in the arm and allows MukB’s hinge to approach the head/MukEF module. Importantly, comparison of SEC profiles from native and cross-linked material suggest that native MukBEF adopts mostly a closed conformation under the conditions used (Fig. 1d).

It has been noted in previous studies that other SMC arms sometimes contain kinks (Anderson et al., 2002; Haering et al., 2002; Hons et al., 2016; Yoshimura et al., 2002). This led us to address whether eukaryotic cohesin would also be able to adopt a defined folded conformation similar to that of MukBEF. We purified budding yeast cohesin containing Smc1, Smc3, the kleisin Scc1 and the HAWK protein Scc3 produced in insect cells, and as for MukBEF, stabilized the complex by mild cross-linking with BS3 and imaged it by negative stain EM (Fig. 1g, Supplementary Fig. 2b, c). The complex appeared in a folded conformation resembling that of MukBEF and reference-free 2D classification revealed well-resolved features in the averages. The head/kleisin/HAWK (Smc1/Smc3 ATPase heads, Scc1, Scc3) module of cohesin is visible as a cherry-shaped density at one end of the complex. It is adjacent to a small constriction that likely represents the conserved head-proximal coiled-coil discontinuity called ‘joint’ (Diebold-Durand et al., 2017; Gligoris et al., 2014). The cohesin hinge, which is larger than the MukB hinge, is visible as a circular density in direct vicinity of the joint. The cohesin elbow is located at an off-center position within the SMC arms in contrast to MukBEF, but, similar to MukBEF, allows them to bend at an angle close to 180 degrees. We conclude that the ability to bend at an SMC elbow is shared by two very distantly related SMC-kleisin complexes.

### Identification of the elbow position in MukBEF and cohesin

To ascertain where the elbow might be located at the sequence level, we used mass-spectrometry to identify the BS3 cross-linked residue pairs in both MukBEF and cohesin. We observed 176 distinct inter-subunit cross-links for MukBEF (Fig. 2a) as well as 352 additional intra-subunit cross-links, whereas analysis of cohesin identified 241 inter- and 503 intra-subunit cross-links (Supplementary Fig. 2d). Using spatial information from crystal structures of the MukEF and the MukB head / MukF C-terminal winged-helix domain (cWHD) subcomplexes, respectively, we computed kernel density estimates for the distribution of inter-subunit cross-links (Fig. 2b) (Fennell-Fezzie et al., 2005; Woo et al., 2009). The distribution of observed cross-links is congruent with the position of known subunit interfaces, indicating that our cross-linking experiment faithfully reports the structure of the complex. We used the same approach to localize regions at the MukB hinge that cross-linked to head-proximal sites and the MukEF module, respectively (Fig. 2c). Cross-links clustered at a coiled-coil region near the hinge (Li et al., 2010), consistent with the idea that the complex folds at an elbow. To pinpoint the elbow region precisely, we next mapped all MukB coiled-coil residues onto a unified sequence coordinate system along the arm (accounting for the antiparallel nature of the SMC arm coiled coils), using available disulfide cross-linking data as a guide (Weitzel et al., 2011) (Fig. 2d). We then filtered intramolecular cross-links in MukB for long-distance residue pairs in this coordinate system and determined the midpoint for each pair. If the coiled-coil arm folds at a defined elbow position, then the midpoints should reveal it, and indeed, midpoints clustered at a central region of the MukB arm (Fig. 1d). As a negative control, clustering was not observed in randomly permutated data (Supplementary Fig. 2e). Kernel density estimates produced a pronounced peak close to the 180th coiled-coil residue in the arm coordinate system (corresponding to MukB residues 427 and 970 on the N- and C-terminal coiled-coil strands, respectively).

**Figure 2.**
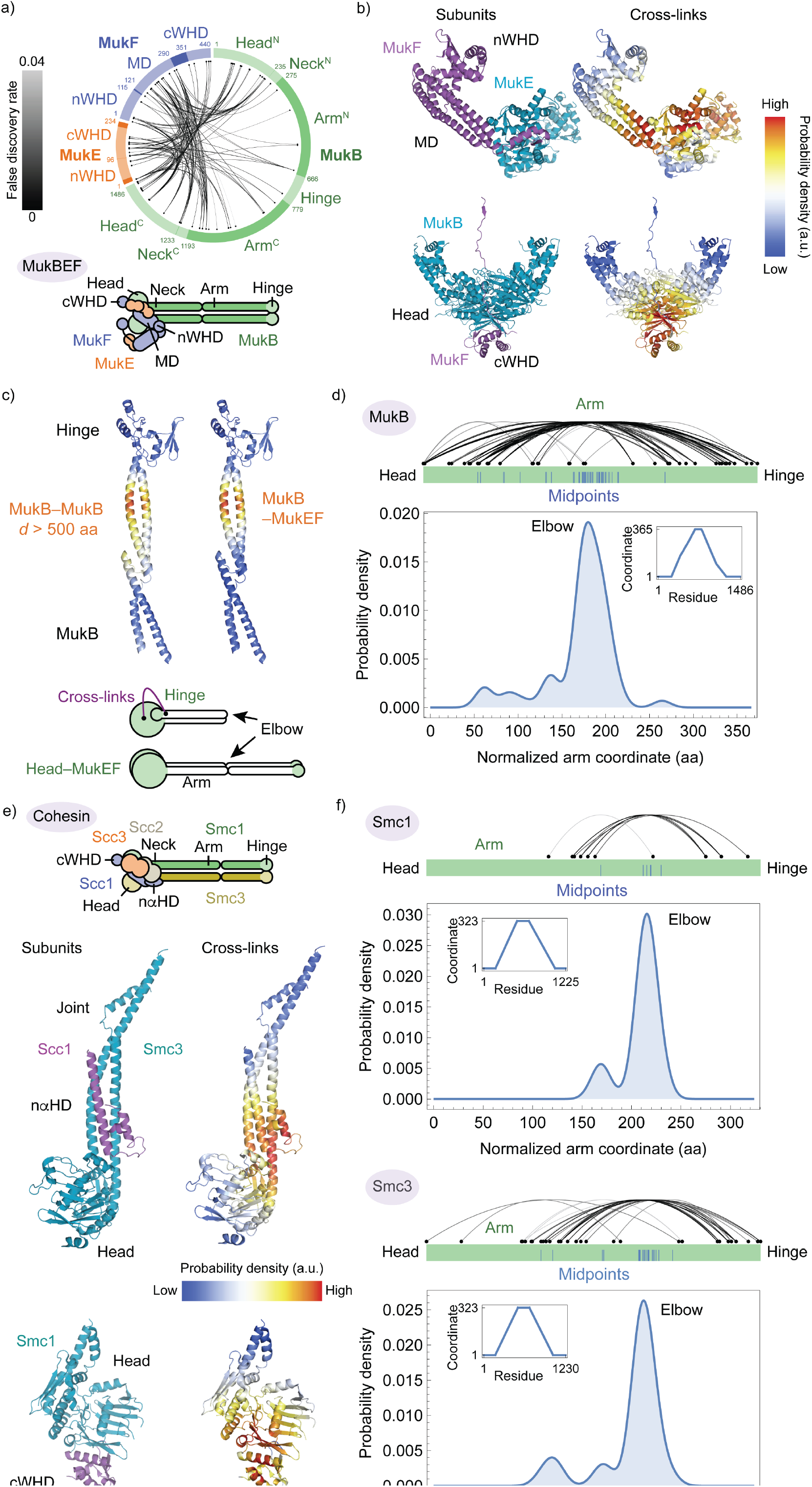
Elbow positions revealed by cross-linking/mass spectrometry. **(a)** Inter-subunit cross-links of MukBEF. Links are colored according to their false discovery rate (FDR). The bottom panel illustrates a likely topology of the complex. **(b)** Kernel density estimates for the position of cross-link sites mapped onto MukBEF subcomplex structures. Cross-link probability densities were mapped onto the *E. coli* MukEF subcomplex (PDB: 3EUH, top) and the partial structure of the *H. ducreyi* MukBEF head module (PDB: 3EUH, bottom). **(c)** Kernel density estimates for long-distance cross-links at the MukB hinge. Probability density for MukB cross-links to MukB sites located at least 500 aa away (left) or to MukEF (right). The bottom panel illustrates an explanation for the observed cross-linking pattern. **(d)** Identification of the MukB elbow region. Long-distance cross-links in a coordinate system along the coiled-coil arm and their midpoints are shown on top. The bottom panel shows the kernel density estimate for the midpoint positions. An inset shows the piecewise interpolation function used to map residue numbers to the arm coordinate system. **(e)** Kernel density estimates for the position of cross-link sites mapped onto cohesin subcomplex structures (PDB: 4UX3, top; 1W1W, bottom). The top panel illustrates the topology of the complex. **(f)** Identification of the cohesin elbow region as in (d).

We used a similar approach to identify the elbow’s position in a cohesin complex comprising Smc1, Smc3, Scc1, Scc3 and the loader protein Scc2. As was the case for MukBEF, kernel density estimates for inter-subunit cross-links are in good agreement with available crystal structures (Fig. 2e) (Gligoris et al., 2014; Haering et al., 2004). Consistent with our observations by EM, the arms of Smc1 and Smc3 both showed midpoint clustering of cross-links at a position away from the center, indicating the presence of the elbow close to residues 391/806 in Smc1 (coiled-coil residue 215 in the arm coordinate system) and 396/808 in Smc3 (residue 212 in the arm coordinate system), respectively. These findings suggest that cohesin’s elbow is shifted towards the hinge, in contrast to MukBEF’s center position (Fig. 2f, Fig. 1c, f, g). Using the same method, we re-analyzed published cross-link/mass-spectrometry (CLMS) data for human and budding yeast cohesin (Chao et al., 2017; Huis in ‘t Veld et al., 2014) and obtained similar results (Supplementary Fig. 2f). We conclude that although cohesin and MukBEF each contain a defined elbow that enables folding, its relative position within the SMC proteins appears to be different.

### Structure of the MukB elbow

To investigate if the MukB arm contains structural features that would allow it to bend at the elbow position, we purified a fusion construct between matching N- and C-terminal fragments containing the elbow as determined above and solved its structure by X-ray crystallography (Fig. 3a). Consistent with findings from disulfide cross-linking experiments, the arm contains two coiled-coil discontinuities or “knuckles” in this region (Weitzel et al., 2011). The knuckle that has been previously named K1/2 is at a central position, joining the coiled-coil regions formed by helices α1/α7 and α2/α6. Knuckle K1/2 is followed by the K2/3a break formed by helices α3, α4 and α5. Mapping the long-distance cross-link midpoints onto the structure identifies the K1/2 break as the elbow (Fig. 3b, 2d). In the crystal, the elbow adopts an extended and gently bent conformation. It contains an “anchor” segment in its a1 helix, which is part the N-terminal coiled-coil region (Fig. 3c). This anchor helix connects to α2 via a loop. The C-terminal part of the coiled-coil segment winds around the elbow anchor helix, starting at α6 that connects to α7 via a distorted helical stretch. A conserved Tyr residue (Y416) (Fig. 3b, c, Supplementary Fig. 3) is wedged into the α6/α7 connection and might contribute to its distortion by obstruction with the bulky Tyr sidechain. The tip of α6 in the C-terminal coiled-coil region forms a short interface with the anchor helix of the N-terminal coil (Fig. 3c). It is conceivable that unzipping of this interface might further destabilize the α6/α7 connection to allow the elbow to bend and fold. Codon substitutions at the chromosomal *mukB* locus that either changed Leu960, located at the α1/ α6/anchor helix interface, to Glu (L960E) or changed the central Tyr416 to Asp or Pro (Y416D and Y416P, respectively) caused a *mukB* null phenotype: mutant strains were not viable on rich media at 37 °C, despite MukB proteins being present at wild-type levels **(Supplementary Fig. 3**). These finding suggest that an intact elbow region is critical for MukBEF activity *in vivo*. Although the structural details of the folded conformation remain to be determined, our findings support the notion that bending of MukBEF occurs at a predetermined, structurally defined and essential coiled-coil discontinuity in its SMC arms.

**Figure 3.**
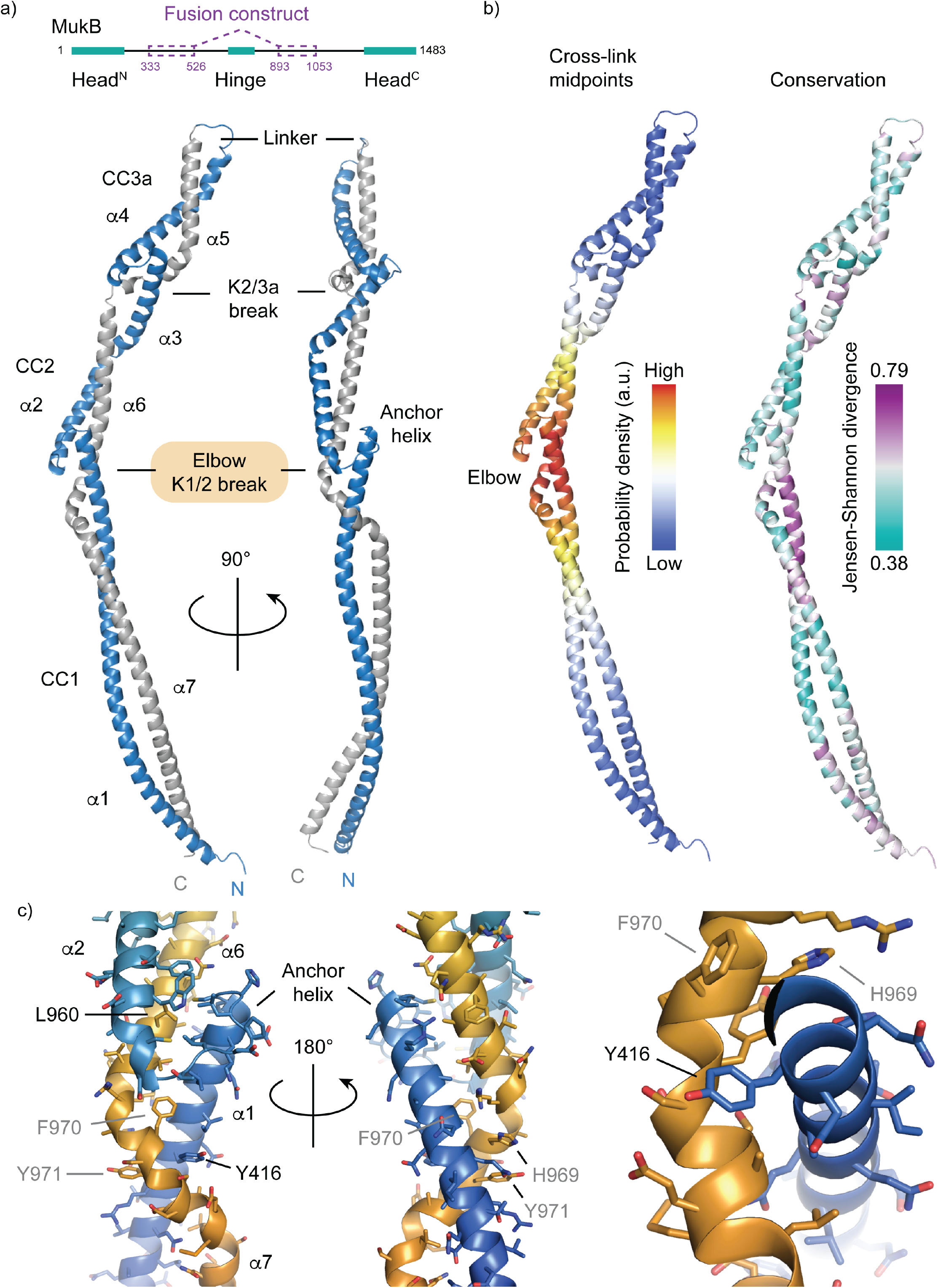
Structure of the MukB elbow. . **(a)** Crystal structure of an *E. coli* MukB arm fragment. The top panel illustrates the design of the fusion construct used for crystallography. The bottom panel shows the refined atomic model obtained from the X-ray diffraction experiment. **(b)** Identification of the elbow. Cross-link midpoint density (see Fig. 2d) was mapped onto the structure (left). The right panel shows sequence conservation (Jensen-Shannon divergence) mapped onto the structure (high conservation: purple, low conservation: cyan). **(c)** Structure of the elbow. The C-terminal coiled-coil helix is distorted (kinked) close to the conserved Tyr416 on the N-terminal helix. Residues for visual reference between the views are shown in grey. Residues targeted by mutagenesis (Supplementary Fig. 3) are highlighted in black.

### Proximity of cohesin HAWK Pds5 and the hinge *in vivo*

A crucial question is whether the coiled coils of SMC-kleisin complexes adopt a folded conformation *in vivo* as well as *in vitro*. We reasoned that if such folding occurred at cohesin’s elbow, then proximity of its hinge domain to ATPase head proximal sequences might permit site-specific chemical cross-linking between residues within the hinge and those associated with ATPase heads. To this end, we generated yeast strains in which residues within the Smc1 hinge were substituted by the unnatural amino acid BPA (p-benzoyl L-phenylalanine) **(Fig. 4a**), the sidechain of which can be activated by ultra-violet (UV) light to cross-link to residues in its vicinity. After UV treatment of intact cells, we immunoprecipitated cohesin and analyzed the cross-linking reaction by Western blotting (Fig. 4b). An Smc1 mutant with a BPA substitution at Glu593 efficiently cross-linked to Smc3 (Fig. 4b) because this residue is positioned directly at the Smc1/Smc3 hinge interface. Strikingly, a BPA substitution mutant of Lys620, located on the coiled-coil distal face of the hinge, efficiently cross-linked to a large protein other than Smc3. This protein was identified as Pds5 (Fig. 4b). We verified that BPA was incorporated into Smc1(K620BPA) (Supplementary Fig. 4) and that cross-linking between Smc1(K620BPA) and Pds5 was dependent on UV treatment (Fig. 4c). Pds5 is a HAWK protein that binds Scc1 sequences close to the kleisin’s N-terminal domain associated with the coiled coil emerging from the Smc3 ATPase head (Lee et al., 2016; Muir et al., 2016). Importantly, mutation of the Pds5 binding site in Scc1 by substitution of Val137 for lysine (V137K) greatly diminished Pds5 recruitment and prevented cross-linking to the Smc1(K620BPA) hinge (Fig. 4d) (Chan et al., 2013).

**Figure 4.**
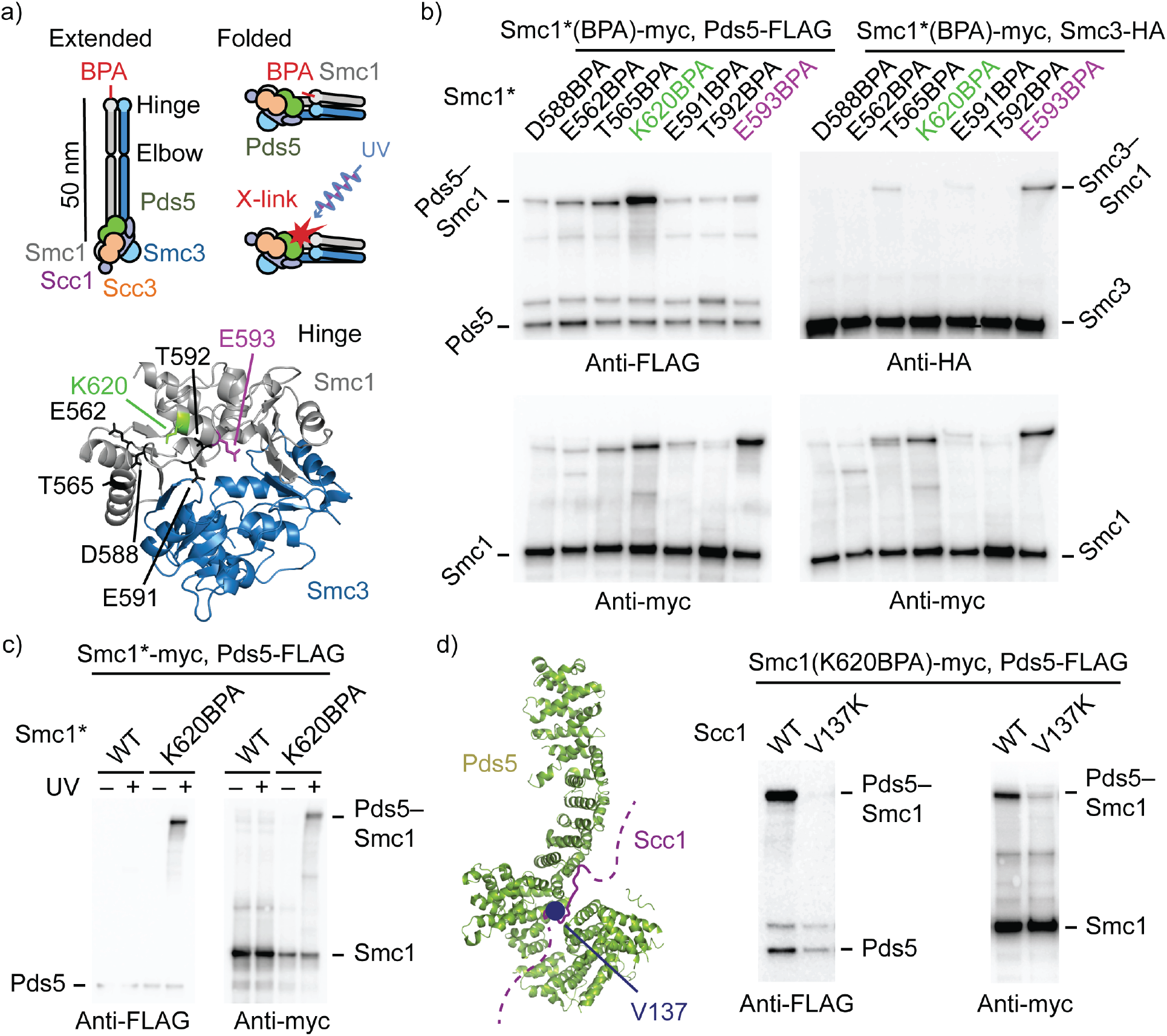
*In vivo* cross-linking of Pds5 to the Smc1 hinge. **(a)** Illustration of the BPA cross-linking experiment (top) and mapping of tested BPA substitutions onto a homology model of the cohesin hinge (bottom). **(b)** Screen for Smc1(BPA) cross-links to Pds5 and Smc3. BPA was incorporated into the indicated Smc1 positions, cells were treated with UV, cohesin was immunoprecipated via a PK9-tag on Scc1 and products were analyzed by Western blotting. (**c**)UV-dependent cross-linking of Smc(K620BPA) and Pds5. Cells were either treated or not treated with UV and products were analyzed as in (b). **(d)** Cross-linking of Smc(K620BPA) and Pds5 depends on Pds5 binding to Scc1. The left panel shows the position of Scc1 V137 in its Pds5 binding site (mapped to the *L. thermotolerans* structure, PDB: 5F0O). The right panel shows a cross-linking experiment of Smc1(K620BPA) in the presence of Scc1(V137K)-PK9 as in (b).

At present, we cannot exclude the possibility that cross-linking between Pds5 and the Smc1 hinge occurs between two different cohesin complexes or that Pds5 binds close to the hinge in a way that only indirectly depends on Scc1. However, if cross-linking happens within a single cohesin complex and Pds5 contacts the hinge while bound to Scc1, then this would necessitate a folded conformation similar to that observed by EM (Fig. 1g). We note that it has previously been observed that a fluorescent tag inserted into the Smc1 hinge of cohesin produces a weak FRET signal when combined with a tag on the head proximal HAWK subunit Pds5 (Mc Intyre et al., 2007), which is consistent with our observations. We conclude that folding of cohesin’s coiled coils most probably occurs *in vivo* as well as *in vitro*.

### Conservation of the SMC elbow

It has been noted before that coiled-coil prediction profiles for SMC sequences often contain at least two considerable drops in coiled-coil probability within both the N- and C-terminal parts of the arm (Waldman et al., 2015). One of the predicted breaks is located close to the SMC heads, and another one is often found at a more central position within the arm. We wished to confirm and corroborate these findings by extending the analysis to a large set of protein sequences. We predicted coiled-coil probabilities for hundreds of individual full-length sequences using MARCOIL (Delorenzi and Speed, 2002), extracted profiles for N- and C-terminal halves, aligned them on the arm center, and averaged the profiles to remove noise **(Fig. 5**). The aggregate profiles for different classes of SMC proteins clearly indicate the position of the head proximal coiled-coil discontinuity, mapping it to the structurally conserved joint (Diebold-Durand et al., 2017; Gligoris et al., 2014). Importantly for our work, the profiles also predict the presence of a centrally located coiled-coil discontinuity in most, if not all, SMC protein families with high confidence as judged by random resampling (Fig. 5). For MukB, the predicted central position is in excellent agreement with the elbow position estimated here experimentally by cross-linking / mass-spectrometry (minimum coiled-coil probability at residues 432 and 970, maximum cross-link midpoint probability density close to residues 427 and 970). These residues are located directly within the K1/2 break present in our crystal structure (Fig. 3). Similarly, the predicted elbow positions for Smc1 and Smc3 (residues 374/790 and 397/796, respectively) are close to our experimental estimates (residues 391/806 and 396/808, respectively). It appears that the prediction method is accurate for the two distantly related SMC-kleisin complexes, which we have investigated here, and hence likely generalizes to other SMC proteins. In addition, the coiled coils of both bacterial and archaeal Smc proteins (*B. subtilis* and *P. yayanosii*) contain a discontinuity close to the elbow position predicted by our aggregate profiling approach (Diebold-Durand et al., 2017; Waldman et al., 2015) (Supplementary Fig. 5). Interestingly, in *B. subtilis* Smc this region is among the few that tolerates peptide insertions (Bürmann et al., 2017). Among Smc proteins, a predicted elbow is particularly apparent in profiles of naturally occurring short variants (Fig. 5). We conclude that a central coiled-coil discontinuity is present in most if not all classes of SMC proteins, indicating that the ability to bend at a defined elbow is likely a fundamental feature.

**Figure 5.**
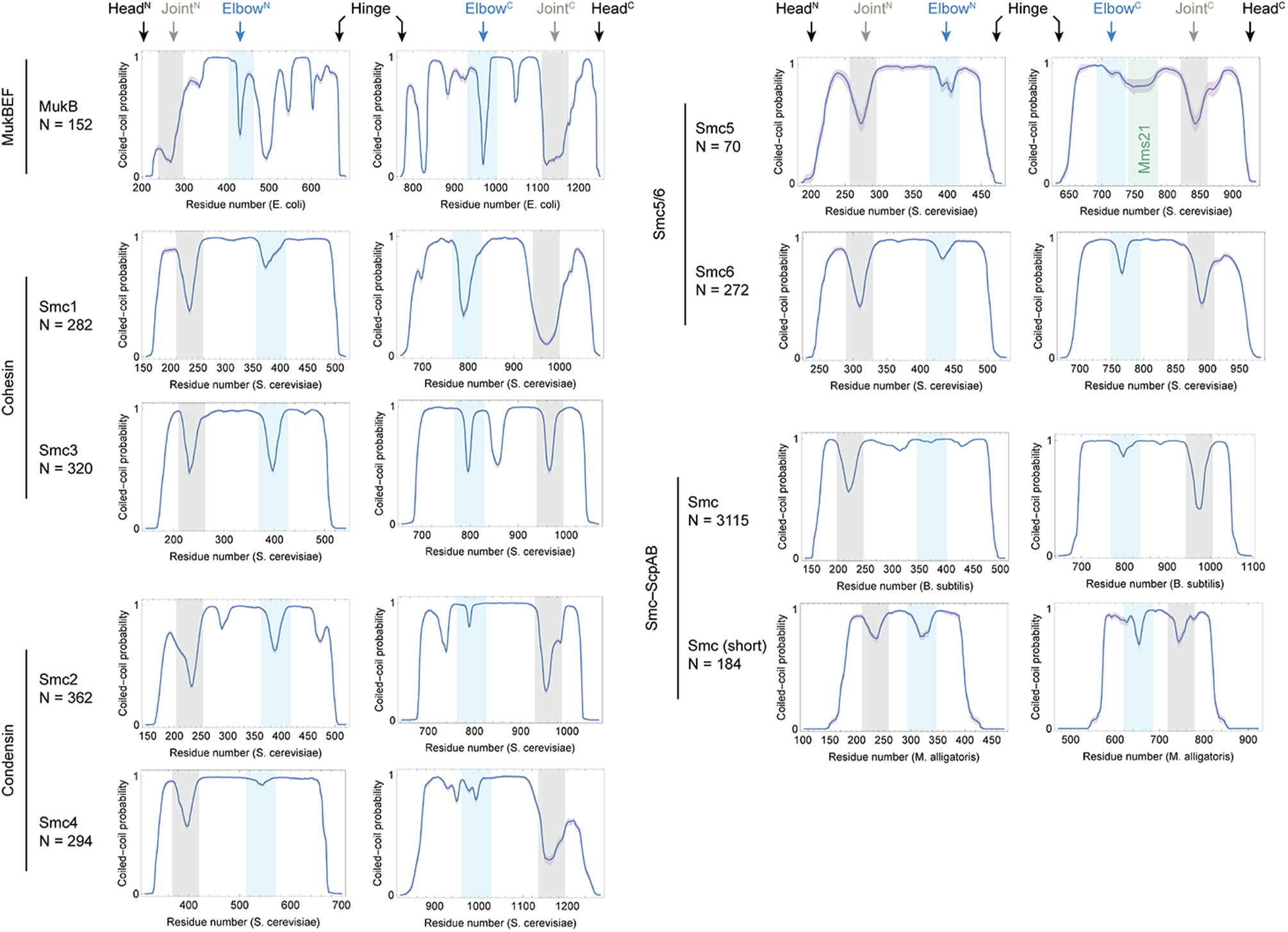
Conservation of the SMC elbow. Coiled-coil prediction profiles for a diverse set of SMC protein sequences were generated by MARCOIL. Profiles for N- and C-terminal parts of the arms were separately aligned on their center coordinate and averaged. 95 % confidence intervals (purple shading) were estimated by 100 times random resampling with replacement. The Mms21 binding site of Smc5 is highlighted in green. N, number of sequences used for generating the respective aggregate profiles.

## Discussion

### Conformational states and their interconversions in SMC-kleisin complexes

The first electron microscopic images of isolated SMC proteins were obtained by rotary metal shadowing of mica-adsorbed proteins and revealed a characteristic shape: positioned at the ends of a long coiled-coil arm were identified two globular domains, a hinge dimerization domain and a head ATPase domain (Haering et al., 2002; Melby et al., 1998; Niki et al., 1992). In these early studies, SMC dimers held together by their hinges were largely observed as V-, I- (rod) or O-shaped particles. Furthermore, it was noticed that the SMC arms would sometimes be kinked (Anderson et al., 2002; Haering et al., 2002; Hons et al., 2016). Other studies, employing atomic force microscopy, have suggested that the isolated SMC2-4 heterodimer of condensin may adopt a compact conformation (Yoshimura et al., 2002) or may have highly flexible arms with a short persistence length (Eeftens et al., 2016). The apparent presence of coiled-coil breaks within SMC arms prompted the prediction that “the coiled coil undergoes a dramatic conformational change to allow a direct interaction between the hinge domain and the head domain or a head-proximal portion of the coiled coil” (Waldman et al., 2015).

Here, we demonstrate that two substantially diverged SMC-kleisin complexes, namely bacterial MukBEF and eukaryotic cohesin, are able to adopt a well-defined folded conformation that brings their hinge into proximity of their heads. Folding of the complexes occurs at a centrally positioned coiled-coil discontinuity, the ‘elbow’, that is present in most if not all SMC proteins. The elbow is apparent as a sharp kink also in cryo-EM images of MukBEF particles, without the use of surface immobilization, dehydration, staining or mechanical probing of the sample (Supplementary Fig. 1), and it is detectable by in-solution cross-linking and mass spectrometry. Observed contact sites are fully consistent with a folded state and are in excellent agreement with computational predictions for the elbow position and also crystallographic data (Figs. 2, 3, 5). Size-exclusion chromatography of MukBEF suggests that a considerable fraction of this complex adopts a folded conformation **(Fig. 1d**). As indicated by SEC and negative stain EM, a smaller fraction adopts an extended (’I’ or rod) conformation, which resembles the shape of *B. subtilis* Smc-ScpAB (Diebold-Durand et al., 2017; Soh et al., 2015). Interestingly, treatment of MukBEF with the cross-linker BS3 strongly enriches the extended rod fraction. Hence, we would like to propose that MukBEF switches between folded and extended forms, and that reaction with BS3 artificially triggers this switch.

If MukBEF and cohesin are able to alternate between folded and extended states, then it is likely that the interconversion is coupled to their DNA binding and ATP hydrolysis cycle. The SMC arms are firmly anchored in the ATPase heads, which are ideally positioned to drive such a conformational change. A link between the ATPase cycle and arm conformation is supported by site-specific cross-linking experiments with Smc-ScpAB that indicate a conformational change in its coiled coil upon binding of ATP and DNA (Minnen et al., 2016; Soh et al., 2015). This conformational change has been interpreted as a disengagement of the arm/arm interface, converting Smc-ScpAB from a rod-like to ring-like state. If both folded and extended conformations interconvert in MukBEF and cohesin, we suspect that they may do so via an intermediate that accommodates considerable strain in its arms. Such an intermediate might correspond to this “open” or ring-like state (Fig. 6a).

**Figure 6.**
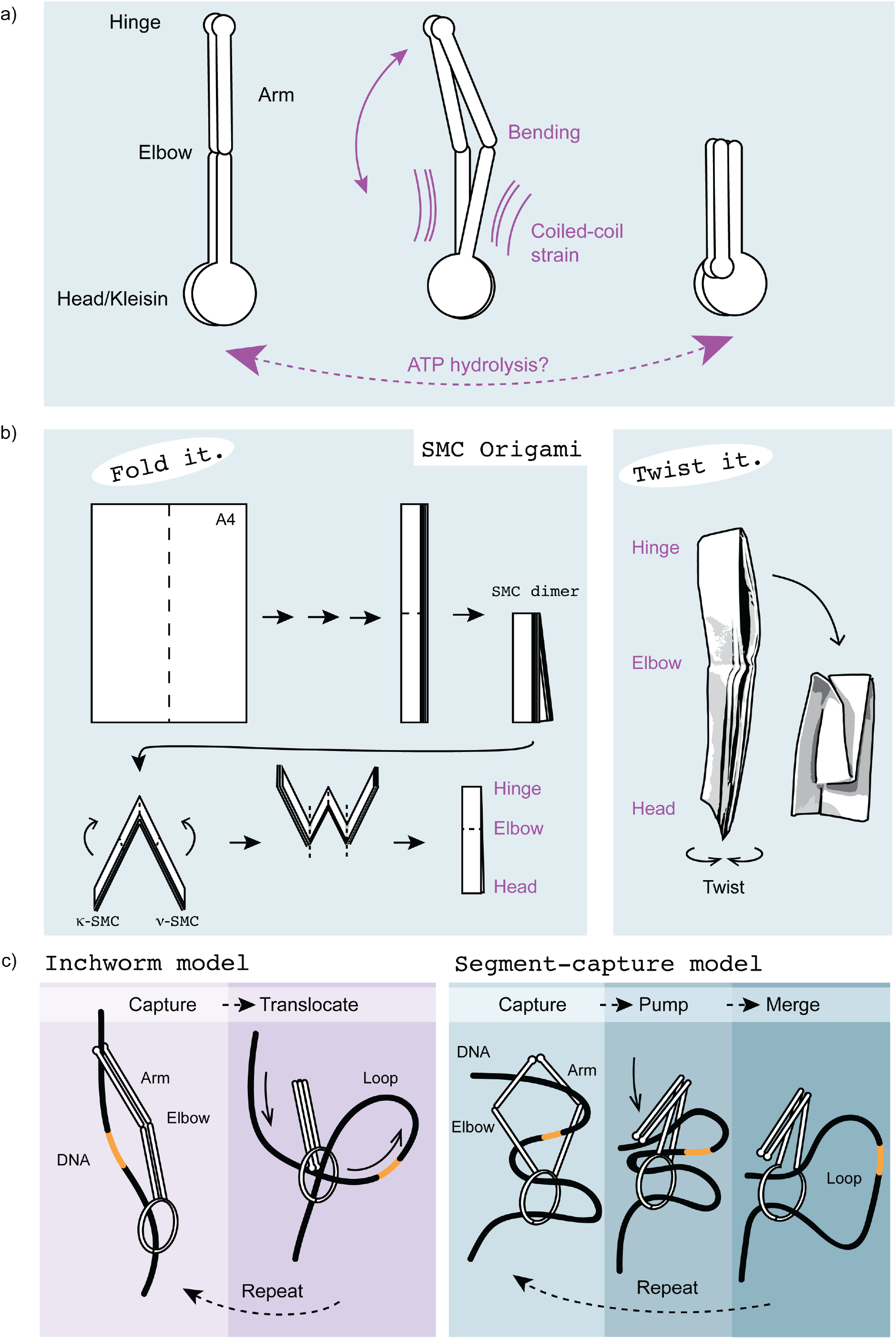
Model for DNA translocation by relative movements of DNA binding sites. **(a)** Model for conformational switching of SMC-kleisin complexes. Transitioning between extended and folded states might be driven by the ATPase cycle introducing mechanical strain into the SMC arms. **(b)** Paper model: conversion between extended and folded states is achieved by twisting the arms of the model. **(c)** Models for DNA translocation and loop extrusion involving a folded state. (left) “Inchworm translocation” using distance changes between two DNA binding sites, one of which might be a topological entrapment device/ring. Folding at the elbow might cause the distance change. (right) Translocation using the segment-capture mechanism that enlarges a loop held in a bottom chamber by merging with a smaller loop captured in a top chamber. Folding at the elbow might drive DNA from top to bottom.

How could an ATPase driven folding/extension cycle be implemented at the structural level? One conundrum is how an SMC dimer with a central 2-fold symmetry axis is able to adopt a folded state such as those observed here. Making an SMC dimer bend to one side severely breaks this symmetry, as the symmetry dictates that the two arms bend to opposite sides if the same bending angle direction is applied to each SMC coiled coil (Supplementary Figure 6). To bend the 2-fold symmetric dimer at the elbow to one side, there are two options: the monomers might bend into opposite directions within their respective body frames, or they might rotate 180 degrees relative to each other and then bend into the same direction. This insight allows the construction of a simple hypothesis: conformational switching between folded and extended states might be achieved by rotating the arms against each other. This would bring the monomer elbows into an orientation that either is or is not compatible with folding at the elbow, depending on the starting conformation. If such a rotation introduced or removed strain, for example by twisting the heads while keeping the hinges fixed, this could actively promote switching. The reader is encouraged to elaborate on these insights by building and twisting the accompanying paper model (Fig. 6b). Of note, asymmetric binding of SMC proteins by the kleisin appears to be a widely conserved feature and might facilitate the asymmetric twisting (Bürmann et al., 2013; Haering et al., 2002; Zawadzka et al., 2018). A mechanism based on a simple mechanical principle such as this would be robust because it could induce bending at arbitrary positions, only depending on the position of the elbow. However, it requires heads and hinge to have a particular relative angular orientation. Consistent with such a geometric constraint, function of Smc-ScpAB appears to be influenced by the superhelical phase-relationship between the ends of its arms (Bürmann et al., 2017) suggesting that ATPase heads and hinges are attached to the coiled-coil arms in fixed and relevant relative orientations.

### Shape transformations for DNA transactions

SMC-kleisin complexes in pro- and eukaryotes appear to share three possibly overlapping activities that are likely central to their biological function: DNA translocation, DNA entrapment and DNA loop extrusion. How these activities are interconnected and how they are biochemically implemented remain exciting and important questions. Understanding the DNA translocation mechanism appears especially relevant, because it is almost certainly required for DNA loop extrusion. Translocation along DNA depends on ATP hydrolysis by SMC proteins (Ganji et al., 2018; Hu et al., 2011; Minnen et al., 2016), and is likely driven by an internal motor activity at least in condensin (Terakawa et al., 2017). The SMC arms likely play a role in the translocation process because mutations in the arms of bacterial Smc-ScpAB largely uncouple ATP hydrolysis from the movement on chromosomes (Bürmann et al., 2017). Moreover, recent findings indicate a dependency of translocation on the SMC hinge in cohesin because mutations in this domain appear to uncouple nucleotide hydrolysis and translocation in a similar fashion (Srinivasan et al., 2018). Apart from DNA translocation, cross-talk between hinge and head module has been suspected to mediate DNA loading of cohesin (Murayama and Uhlmann, 2015; Xu et al., 2018). We envision that a large-scale conformational change in the arms, such as folding at the elbow, is involved in DNA translocation or entrapment by SMC-kleisin complexes.

How might a folded conformation of SMC-kleisin complexes take part in the translocation reaction? One intriguing possibility is that switching between extended and folded conformations might change the distance between two DNA binding sites, possibly located at the ends of the SMC-kleisin rod, thereby permitting an inchworm-like movement along the substrate (Fig. 6c, Supplementary Fig. 7a). DNA binding activity has been reported for the head domains of MukB, the SMC-like Rad50 and Smc-ScpAB (Lammens et al., 2004; Liu et al., 2016; Löwe et al., 2001; Woo et al., 2009). Moreover, isolated hinge domains of cohesin, condensin, Smc5-6 and Smc-ScpAB are also able to bind DNA (Alt et al., 2017; Chiu et al., 2004; Griese et al., 2010; Soh et al., 2015). The MukB hinge, although lacking a strong interaction with DNA (Ku et al., 2010; Kumar et al., 2017), associates with the DNA binding proteins topoisomerase IV and MatP, suggesting that it might at least come into proximity of the substrate (Nolivos et al., 2016; Vos et al., 2013). Translocation via an inchworm-like mechanism would require at least one of the binding sites to act as a “grapple”, i.e. it must have regulated DNA affinity for capture and release of the substrate (Supplementary Fig. 7a). The second site would act as an “anchor”, keeping the complex attached to DNA while the grapple is released. DNA binding at either site might not be purely enthalpic but may involve entropic (steric or even topological) entrapment of the substrate, similar to the sliding clamp of DNA polymerase (Hedglin et al., 2013). In addition, the DNA binding sites could also be located on different complexes that act in concert, as is clearly a possibility for chromosomal MukBEF (Badrinarayanan et al., 2012). If a distance change in DNA binding sites was responsible for translocation of SMC-kleisin complexes, then loop extrusion might be implemented by addition of a third site that stabilizes the loop (Supplementary Fig. 7b). The HAWK subunits Ycg1 and Scc3 of condensin and cohesin, respectively, are part of a DNA binding site that might act as such an anchor (Ganji et al., 2018; Kschonsak et al., 2017; Li et al., 2018).

It has been proposed that DNA translocation of Smc-ScpAB complexes involves the transition from a rod- to a ring-like state, whereby DNA is captured in between the SMC arms to be pushed from a hinge-proximal site towards a head-proximal compartment (Diebold-Durand et al., 2017; Marko et al., 2018; Soh et al., 2015). Although a central coiled-coil discontinuity is present in bacterial/archaeal Smc proteins (Fig. 5, Supplementary Fig. 5), it is currently not clear whether these proteins are able to adopt a folded conformation. Smc proteins are distant relatives of both MukB and cohesin’s Smc1 and Smc3, and folding might have arisen by convergent evolution between the MukBEF and cohesin complexes. In such a scenario, the SMC elbow might have a twofold role: First, it might support bending of the relatively rigid arms into a ring, which in MukBEF might be assisted by secondary coil-coil discontinuities (Fig. 3) (Li et al., 2010; Weitzel et al., 2011). Second, transition to a folded conformation might build on top of this possibly primordial activity to help pushing DNA from one end of the complex to the other (Fig 6c). However, at present it is unclear whether DNA translocation, or loop extrusion, requires entrapment within the tripartite ring. Clarifying the nature of the DNA bound states of SMC-kleisin complexes and tracing the path of associated DNA is now of utmost importance.

In summary, we show that the evolutionarily distant SMC-kleisin complexes MukBEF and cohesin adopt very similar folded conformations by bending at a central coiled-coil discontinuity, the elbow. We provide evidence that the elbow is a general feature of SMC-kleisin complexes and propose it is involved in a conformational switch that drives DNA transactions of all SMC-kleisin complexes.

## Methods

### Purification of MukBEF

Coding sequences for *E. coli* MukF, MukE and MukB (GeneBank IDs: NP_415442.1, NP_415443.2, NP_415444.1) were inserted as a polycistronic expression construct into a pET-28 derived vector using Golden Gate cloning (Engler et al., 2008). MukB was N-terminally fused to budding yeast His6-SUMO. The complex was produced in *E. coli* BL21-Gold(DE3) grown in ZYP-5052 autoinduction medium at 24 °C (Studier, 2005). Purification of the complex was performed at 4 °C. About 15 g cells were resuspended in 90 mL buffer IMAC (50 mM sodium phosphate, 300 mM NaCl, 20 mM imidazole, pH 7.4 @ 4 °C) including RNase A, DNase I and protease inhibitors and lysed in a high-pressure homogenizer at 172 MPa. The lysate was briefly sonicated to reduce viscosity, and cleared by centrifugation for 30 min at 96,000 x g. The extract was incubated with 25 mL NiNTA agarose (Qiagen) for 30 min. The resin was packed into a gravity flow column and washed with 80 mL IMAC buffer followed by 40 mL SENP buffer (10 mM sodium phosphate, 50 mM NaCl, 20 mM imidazole, pH 7.4 @ 4 °C). The resin was resuspended in 25 mL SENP buffer containing 1 mM DTT and 1 mg GST-hSENP1 protease and incubated for 1 h to cleave off the His6-SUMO-tag. The flow-through containing the complex was collected and combined with an additional 12.5 mL wash of the column. The eluate was then loaded onto a 20 mL Heparin HP column (GE Healthcare), washed with 2 column volumes (CV) of buffer HA (10 mM sodium phosphate, 50 mM NaCl, pH 7.4 @ 4 °C) and eluted with a 20 CV gradient into buffer HB (10 mM sodium phosphate, 1 M NaCl, pH 7.4 @ 4 °C). The complex eluted in two peaks, whereby the low salt peak contained a prominent contaminant. The high salt peak fractions (at about 400 mM NaCl) were pooled and diluted with 4 volumes of buffer (10 mM Tris, 70 mM NaCl, pH 8.0 @ 4 °C). The solution was loaded onto a 5 mL Q HP column (GE Healthcare). The column was washed with 2 CV of buffer QA (10 mM Tris, 200 mM NaCl, pH 8.0 @ 4 °C) and eluted with a 20 CV gradient into buffer QB (10 mM Tris, 1 M NaCl, pH 8.0 @ 4 °C). The complex eluted as a single peak at about 450 mM NaCl. Peak fractions were pooled, concentrated to about 10 mg/mL on a Vivaspin 100k ultrafiltration membrane (Sartorius), aliquoted, frozen in liquid nitrogen and stored at -80° C. An aliquot of MukBEF was then thawed and injected into a Superose 6 Increase 3.2/300 column (GE Healthcare) in T200 buffer (10 mM Tris, 200 mM NaCl, pH 8.0 @ 4 °C) to remove aggregates. The monomer fraction appeared stable for several days as judged by SEC but was freshly used for all experiments. MukBEF from *Desulfovermiculus halophilus* (GeneBank IDs: WP_027370798.1, WP_ 027370797.1, WP_ 027370796.1) was produced and purified similar to the *E. coli* complex.

### Purification of MukB and MukEF and reconstitution of MukBEF complexes

MukB was produced and purified similar to the MukBEF holocomplex. MukEF was produced from a polycistronic expression vector with a His6-SUMO-tag on MukE and purified similar to the holocomplex, but omitting the Heparin step and using Sephacryl S200 as the size-exclusion resin. Complexes were reconstituted similar to the protocols from (Petrushenko et al., 2006) at 2 µM MukB_2_ and 4 µM MukE_4_F_2_ in either 10 mM Tris, 40 mM NaCl, 2 mM MgCl_2_, pH 8.0 (singlets, MukBEF^S^) or in 10 mM Tris, 200 mM NaCl, pH 8.0 (doublets, MukBEF^D^). Reactions were run over Superose 6 Increase in the respective reconstitution buffer and peak fractions were re-injected into Superose 6 Increase in T200 buffer.

### Purification of cohesin

Cohesin expression constructs were cloned as described previously (Petela et al., 2018). Briefly, coding sequences for Smc1, Smc3, Scc1, Scc2 and Scc3 from *S. cerevisiae* (Genbank IDs: NP_116647.1, NP_012461.1, NP_010281.1, NP_010466.3, NP_012238.1) were synthesized with codon optimization for insect cell expression (Genscript). Sequences were individually cloned as Smc1, 8xHis-Smc3, Scc1-2xStrepII, 2xStrepII-Scc3 and 2xStrepII-(151-1493)Scc2 into Multibac vectors (Bieniossek et al., 2008), yielding Smc1-pACEbac1, 8xHis-Smc3-pACEbac1, 2xStrepII-∆N150-Scc2-pACEbac1, 2xStrepII-Scc3-pACEbac1 and Scc1-2xStrepII-pIDC. Tagged constructs contained an HRV 3C protease site (LEVLFQ/GP) in the tag linker. The vectors Smc1-pACEbac1, 8xHis-Smc3-pACEbac1 and Scc1-2xStrepII-pIDC were combined through Gibson assembly and *in vitro* Cre-*lox* recombination yielding a transfer vector for the Smc1-Smc3-Scc1 trimer. The trimer, 2xStrepII-∆N150-Scc2-pACEbac1 and 2xStrepII-Scc3-pACEbac1 transfer vectors were individually transformed into chemically competent DH10EmbacY cells (Vijayachandran et al., 2011). The purified bacmids were transfected into Sf9 cells using Fugene HD reagent (Promega), and the generated P1 viruses were infected into fresh Sf9 cells. The cells were grown in Insect XPRESS protein free medium with L-glutamate (Lonza) at 27 °C for ~72 h, and the harvested cells were frozen in liquid nitrogen.

The frozen pellets of Sf9 culture were re-suspended in lysis buffer (20 mM Hepes pH 7.5, 125 mM NaCl, 1 mM TCEP, and 10 % (w/v) glycerol) supplemented with DNase, RNase, 1 mM PMSF and EDTA-free protease inhibitor (cOmplete, Roche). Cells were lysed by sonication, and the lysates were clarified by ultracentrifugation at 200,000 x g. The clarified lysates were applied to Strep resin (5 mL StrepTrap, GE Healthcare) and eluted with 2 mM desthiobiotin in lysis buffer. 3C protease was added to the eluents to cleave the affinity tags and the cleavage products were further purified by anion exchange columns (HiTrap Q FF or mini Q (GE healthcare)) with buffers of QA (50 mM Tris, 100 mM NaCl, 1 mM TCEP, and 5 % (w/v) glycerol, pH 8.0) and QB (50 mM Tris, 1 M NaCl, 1 mM TCEP, and 5 % (w/v) glycerol, pH 8.0). The fractions were pooled, concentrated using a Vivaspin 100 kDa ultrafiltration membrane (Sartorius). The purified trimer, N∆150-Scc2, Scc3 proteins were then frozen in liquid nitrogen and stored at -80 °C until further use.

### BS3 cross-linking / SEC

An aliquot of MukBEF Q eluate was thawed and injected into a Superose 6 Increase 3.2/300 column in P200 buffer (10 mM sodium phosphate, 200 mM NaCl, pH 7.4 @ 4 °C). The monomer fraction was incubated for 2 h on ice at 0.4 mg/mL with or without 1 mM BS3 and was injected into a Superose 6 Increase 3.2/300 column in T200 buffer (10 mM Tris, 200 mM NaCl, pH 8 @ 4 °C). Chromatography was performed at a flow rate of 40 µL/min.

### Negative stain electron microscopy

For imaging of native MukBEF, an aliquot of Q eluate was thawed and injected into a Superose 6 Increase 3.2/300 column in T200 buffer. The monomer fraction was reinjected, and the peak fraction applied to freshly glow-discharged EMS Cu Mesh 400 continuous carbon grids. Grids were stained with 2 % uranyl acetate and imaged in a Tecnai Spirit microscope (FEI) using an Orius CCD camera at a pixel size of 3.5 Å and an electron dose of 30 e^-^/Å^2^ at 120 kV. For data collection, native MukBEF was applied to Quantifoil CuRh R2/2 Mesh 200 grids covered with a homemade continuous carbon film. The grids were stained with 2 % uranyl formate and imaged on a Tecnai F30 Polara microscope (FEI) with a Falcon III detector using a pixel size of 1.72 Å and an electron dose of 30 e^-^/Å^2^ at 300 kV.

BS3 cross-linked MukBEF was prepared as described above and imaged on EMS Cu Mesh 400 continuous carbon grids stained with 2 % uranyl formate. Data for SEC peak 1 (extended conformation) were collected on a Tecnai Spirit with an UltraScan CCD camera using a pixel size of 3.95 Å and an electron dose of 30 e^-^/Å^2^ at 120 kV. Data for SEC peak 2 (folded conformation) were collected on a Tecnai G2 F20 microscope (FEI) with a Falcon II detector using a pixel size of 2.08 Å and an electron dose of 30 e^-^/Å^2^ at 200 kV.

For imaging of cohesin, the purified trimer and Scc3 were mixed at a 1:1.5 molar ratio and injected into a Superose 6 Increase 3.2/300 column in P200 buffer. The tetramer fraction was incubated with 1 mM BS3 for 2 h on ice and injected into a Superose 6 Increase 3.2/300 column in T200 buffer. Peak fractions were applied to Quantifoil CuRh R2/2 Mesh 200 grids covered with a homemade continuous carbon film and stained with 2 % uranyl formate. Data were collected on a Tecnai G2 F20 microscope (FEI) with a Falcon II detector using a pixel size of 2.08 Å and an electron dose of 30 e-/Å^2^ at 200 kV.

### Cryo-electron microscopy

*D. halophilus* MukBEF at 0.2 mg/mL was applied to glow-discharged Quantifoil CuRh R2/2 Mesh 200 grids, blotted on a Vitrobot (FEI) and plunge frozen in liquid ethane. Particles were imaged on a FEI Titan Krios equipped with a Volta phase plate (Danev et al., 2014) and a Falcon III detector operating in linear mode, using a pixel size of 1.07 Å, defocus of -0.6 µm to -0.8 µm and a total electron dose of 100 e-/Å^2^ at 300 kV.

### Image processing

The contrast transfer function (CTF) for electron micrographs was estimated with CTFFIND-4.1 (Rohou and Grigorieff, 2015). For movies collected on a Falcon III detector, motion correction was performed with Motioncor2 (Zheng et al., 2017). Particle picking and reference-free 2D classification were performed in RELION2 (Fernandez-Leiro and Scheres, 2017).

### Cross-linking/mass spectrometry

For cross-linking/mass spectrometry analysis of MukBEF, aliquots of Q eluate were thawed and injected into a Superose 6 Increase 3.2/300 column in buffer XL (20 mM Hepes, 150 mM NaCl, 5 mM MgCl_2_, pH 7.8 @ 23 °C). The monomer fractions were pooled and incubated at 0.4 mg/mL with 2.5 mM BS3 for 2 h on ice before quenching with 20 mM ammonium bicarbonate for 30 min on ice. The sample was incubated for 2 min at 98 °C in the presence of LDS-PAGE sample buffer (Life Technologies) containing 6 % 2-mercaptoethanol. Reaction products were separated on Criterion TGX 4-20 % SDS-PAGE gels (BioRad).

For analysis of cohesin, the purified trimer, N∆150-Scc2 and Scc3 were mixed at a 1:1.5:1.5 ratio and into a Superose 6 Increase 3.2/300 column in buffer (20 mM Hepes, 150 mM NaCl and 1 mM TCEP, pH 7.7). Pentamer fractions were incubated at 2 mg/mL with 5 mM BS3 for 2 hr at 4 °C and then the reaction was quenched with 50 mM ammonium bicarbonate for 45 min on ice. Reaction products were separated on a Criterion TGX 4-15% SDS-PAGE gel (BioRad).

Gel bands corresponding to the cross-linked species were excised and digested with trypsin (Pierce, Germany) (Shevchenko et al., 2006). The resulting tryptic peptides were extracted and desalted using C18 StageTips (Rappsilber et al., 2003).

For MukBEF, peptides eluted from StageTips were dried in a Vacuum Concentrator (Eppendorf, Germany) and dissolved in running buffer A prior to strong cation exchange chromatography (100 x 2.1 mm Poly Sulfoethyl A column; Poly LC, Colombia, MD, USA). Mobile phases A consisted of 30 % acetonitrile (v/v), 10 mM KH_2_PO_4_ at pH 3, and mobile phase B additionally contained 1 M KCl. The separation of the digest used a non-linear gradient (Chen et al., 2010) at a flow rate of 200 µl/min. Five fractions a 2 min in the high-salt range were collected and cleaned by StageTips for subsequent LC-MS/MS analysis. For cohesin, peptides were fractionated on an ÄKTA Pure system (GE Healthcare) using a Superdex Peptide 3.2/300 (GE Healthcare) at a flow rate of 10 µL/min using 30% (v/v) acetonitrile and 0.1% (v/v) trifluoroacetic acid as mobile phase. Five 50 µl fractions were collected and dried.

Samples for analysis were resuspended in 0.1% v/v formic acid 1.6% v/v acetonitrile. LC-MS/MS analysis was conducted in duplicate for SEC fractions and triplicate for SCX fractions, performed on an Orbitrap Fusion Lumos Tribrid mass spectrometer (Thermo Fisher Scientific, Germany) coupled on-line with an Ultimate 3000 RSLCnano system (Dionex, Thermo Fisher Scientific, Germany). The sample was separated and ionized by a 50 cm EASY-Spray column (Thermo Fisher Scientific). Mobile phase A consisted of 0.1% (v/v) formic acid and mobile phase B of 80% v/v acetonitrile with 0.1% v/v formic acid. Flow-rate of 0.3 μL/min using gradients optimized for each chromatographic fraction from offline fractionation ranging from 2% mobile phase B to 45% mobile phase B over 90 min, followed by a linear increase to 55% and 95% mobile phase B in 2.5 min, respectively. The MS data was acquired in data-dependent mode using the top-speed setting with a three second cycle time. For every cycle, the full scan mass spectrum was recorded in the Orbitrap at a resolution of 120,000 in the range of 400 to 1,600 m/z. Ions with a precursor charge state between 3+ and 6+ were isolated and fragmented. Fragmentation by Higher-energy collisional dissociation (HCD) employed a decision tree logic with optimized collision energies (Kolbowski et al., 2017). The fragmentation spectra were then recorded in the Orbitrap with a resolution of 30,000. Dynamic exclusion was enabled with single repeat count and 60-second exclusion duration.

The fragment spectra peak lists were generated from the raw mass spectrometric data using msConvert (version 3.0.11729) (Chambers et al., 2012) with default settings. A recalibration of the precursor m/z was conducted based on high-confidence (<1% FDR) linear peptide identifications, using an in-house script (Lenz et al., 2018). The recalibrated peak lists were searched against the sequences and the reversed sequences (as decoys) of cross-linked peptides using the Xi software suite (version 1.6.739) (Giese et al., 2016) (https://github.com/Rappsilber-Laboratory/XiSearch) for identification. The following parameters were applied for the search: MS1 accuracy = 3 ppm; MS2 accuracy = 10 ppm; enzyme = trypsin (with full tryptic specificity) allowing up to four missed cleavages; cross-linker = BS3 with an assumed reaction specificity for lysine, serine, threonine, tyrosine and protein N termini); fixed modifications = carbamidomethylation on cysteine; variable modifications = oxidation on methionine, hydrolysed / aminolysed BS3 from reaction with ammonia or water on a free cross-linker end. The identified candidates were filtered to 5% FDR on link level using XiFDR (Fischer and Rappsilber, 2017).

### Analysis of cross-linked residue pairs

For mapping of contact sites, kernel density estimation was performed on a per-protein basis for the C-alpha coordinates of cross-link residue pairs present in the respective structures. The coordinates were convolved with a three-dimensional Gaussian kernel (bandwidth: 25 Å), and the resulting probability density distributions were sampled at all C-alpha coordinates of the respective proteins.

For the determination of long-distance cross-link midpoints, we first mapped each residue onto a unified coordinate system along the arm (running from the head at coordinate 0 to the hinge at coordinate 1). Using this approach, residues that are at the same position along the coiled-coil axis but reside on opposite coiled-coil helices map to the same coordinate. For MukB, we used the coiled-coil register established by disulfide cross-linking to build a piecewise linear interpolation function for the coordinate transformation (Weitzel et al., 2011). For each arm segment with a length mismatch between N- and C-terminal parts we used the shorter part as the length of the segment. Residues located in the head were mapped to coordinate 0, residues in the hinge were mapped to 1, and residues located on either the N- or C-terminal arm helix were mapped to the interval (0, 1) according to the disulfide cross-linking data. We used the same approach for cohesin but with single interval interpolation for the N- and C-terminal helices, respectively, due to the mostly unknown coiled-coil register. Finally, coordinates were scaled to an arm length in amino acids (aa) given by the sum of the individual arm segments (MukB: 365 aa, Smc1/3: 323 aa). Cross-link residue pairs with coordinates transformed according to this procedure were filtered for distances of at least 100 aa, and the corresponding midpoints were determined. Kernel density estimation for the distribution of midpoints was performed by convolution with a Gaussian kernel (bandwidth: 10 aa). Cross-link data are available in **Table S1**.

### Purification of the MukB elbow fragment

Residues 333-526 of MukB (GenBank ID: NP_415444.1) were fused to residues 893-1053 using an SGGS linker. The construct contained a C-terminal GSHHHHHH tag and was inserted into a pET-16 derived vector using Golden Gate cloning (Engler et al., 2008). Selenomethionine (SeMet) labeled protein was produced in *E. coli* BL21-Gold(DE3) grown in autoinduction medium PASM-5052 at 24 °C (Studier, 2005). Purification was performed at 4 °C. About 40 g of cells were resuspended in 200 mL buffer NA (50 mM sodium phosphate, 300 mM NaCl, 40 mM imidazole, 1 mM DTT, pH 7.4 @ 4 °C) containing DNase I, RNase A and protease inhibitors. Cells were lysed in a high-pressure homogenizer at 172 MPa, the lysate was briefly sonicated to reduce viscosity, and was cleared by centrifugation at 96,000 x g for 30 min. The extract was passed over a 5 mL HisTrap HP column (GE Healthcare), the column was washed in 10 CV NA and eluted with buffer NB (40 mM sodium phosphate, 240 mM NaCl, 400 mM imidazole, 1 mM DTT, pH 7.4 @ 4 °C). The eluate was diluted in 2 volumes of buffer (10 mM Tris, 1 mM TCEP, pH 8.0 @ 4 °C) and loaded onto a 5 mL Q HP column (GE Healthcare). The column was washed with 3 CV of buffer QA (10 mM Tris, 100 mM NaCl, 1 mM TCEP, pH 8.0 @ 4 °C) and eluted with a 20 CV gradient into buffer QB (10 mM Tris, 1 M NaCl, 1 mM TCEP, pH 8.0 @ 4 °C). Peak fractions were pooled and concentrated in a Vivaspin 10k filter (Sartorius) to about 10 mL before injection into a Sephacryl S200 26/60 column (GE Healthcare) in buffer SEC (10 mM Tris, 150 mM NaCl, 1 mM EDTA, 1 mM TCEP, 1 mM NaN3, pH 7.4 @ 23 °C). Peak fractions were pooled, concentrated to 21 mg/mL in a Vivaspin 10k filter, aliquoted, frozen in liquid nitrogen and stored at -80 °C. The construct had lost its N-terminal methionine as judged by ESI-TOF mass-spectrometry.

### Crystallization of the MukB elbow and structure determination

An aliquot of the MukB elbow construct was thawed and exchanged into buffer X (10 mM Mes, 150 mM NaCl, 1 mM EDTA, 1 mM TCEP, 1 mM NaN3, pH 6.5 @ 23 °C) using a Zeba Spin column (Thermo Scientific). Crystallization conditions were found by screening a set of 1728 conditions using an in-house robotic setup (Stock et al., 2005). Crystals grew as thin plates at 19 °C in sitting drops with mother liquor ML1 (22 % PEG 3350, 0.25 M sodium thiocyanate) or mother liquor ML2 (23.5 % PEG 3350, 2 % PEG 4000, 0.375 M sodium thiocyanate). Crystals mounted in nylon loops were dipped into cryoprotectant solution (23 % PEG 3350, 0.257 M sodium thiocyanate, 30 % glycerol in buffer X) before freezing in liquid nitrogen. X-ray diffraction data were collected at Diamond Light Source I04-1 at a wavelength of 0.91587 Å. Several crystals were tested, whereby a crystal grown in ML1 diffracted to the highest resolution (2.6 Å) but produced weak anomalous signal. A crystal grown in ML2 diffracted to about 3.0 Å but yielded good anomalous signal. The space group of the crystals was determined as P2_1_ using Pointless (Evans and Murshudov, 2013). Diffraction data were integrated with XDS, scaled and merged with Aimless, and converted to structure factor amplitudes with Ctruncate (Evans and Murshudov, 2013; Kabsch, 2010). Automated structure solution with CRANK2 using data from the ML2 crystal yielded an almost complete initial model (Winn et al., 2011). This was used as a search model for molecular replacement in Phaser AutoMR with the ML1 dataset (Bunkoczi et al., 2013). A random set of 5 % of the reflections was retained for validation, and the model was rebuilt from scratch using Buccaneer (Cowtan, 2006). The model was iteratively refined by manual building in Coot and reciprocal space refinement using REFMAC5 (Emsley and Cowtan, 2004; Murshudov et al., 2011). At later stages, manual building was alternated with reciprocal and real space refinement using Phenix.refine (Afonine et al., 2012). Data collection and refinement statistics are listed in Table S2.

### *E. coli* strain construction and growth

*E. coli* strains are based on MG1655 (DSM 18039). All chromosomal modifications were done by λ-Red recombineering using a temperature sensitive plasmid carrying the λ phage genes *exo*, *bet* and *gam* under control of the heat-labile CI857 repressor (Datta et al., 2006). A *neoR* coding sequence was joined with a terminator sequence by Golden Gate assembly and the product was integrated downstream of the *mukFEB* terminator. An in-frame deletion of *mukB* and a *mukB-HaloTag* allele were constructed similarly, terminated by the *mukFEB* terminator and linked to the *neoR* cassette downstream of the operon. For construction of marker-free strains carrying point mutations in *mukB*, target regions were first replaced by a cassette containing the counter-selection marker *pheS(R251A, A294G*) (Miyazaki, 2018) linked to a *hygR* selection marker. The cassette was then ejected by recombination with a PCR product containing the point mutation and counter-selection on media containing 2.5 mM 4-chlorophenylalanine. Strains with a *mukB* null phenotype were grown on LB or TYE at 22 °C or on M9 (lacking thiamine) at 37 °C. Recombineering plasmids were cured in either LB at 37 °C (functional *mukB* alleles) or in M9 (lacking thiamine) at 37 °C (*mukB* null alleles). Strains were single-colony purified and verified by marker analysis, PCR and Sanger sequencing. Strains are listed in Table S3. Phenotypic analysis was performed by streaking on TYE and growth at 37 °C for 13 h.

### *E. coli* HaloTag labelling

Cells were grown to stationary phase in LB at 22 °C, diluted in LB to OD_600_ = 0.02, and grown to OD_600_ = 0.3-0.4 at 37 °C (non-permissive temperature). Cultures were mixed with 30 % (w/v) ice and harvested by centrifugation. Cells were resuspended in B-PER (Thermo Fisher) containing 1 mM EDTA (pH 7.4), 5 µM HaloTag-TMR substrate (Promega), Ready-Lyse lysozyme (Epicentre), Benzonase (Sigma), protease inhibitor cocktail (Roche) and 28 mM 2-mercaptoethanol. Samples were incubated for 10 min at 37 °C, mixed with LDS sample buffer (Thermo Fisher), incubated at 95 °C for 5 min and resolved by SDS-PAGE. Gels were scanned on a Typhoon imager (GE Healthcare) using a Cy3 filter setup, and subsequently stained with InstantBlue (Expedeon).

### Yeast strain construction

Smc1-myc9 with its endogenous promoter was cloned into the LEU2 2μ plasmid YEplac181 and the codon for E620 was replaced with the amber codon TAG. The TRP1 2μ pBH61 expressing the *E. coli* nonsense suppressor tRNA/tRNA synthetase system was a gift from Steven Hahn’s lab. The endogenous Scc1 and Pds5 were fused to 9xPK and 6xFLAG epitope tags at their C-terminus, respectively. All strains are derived from the W303 background and are listed in Table S3.

### *In vivo* photo cross-linking

The yeast stains bearing the TAG-substituted Smc1-myc9 plasmid and pBH61 were grown in −Trp −Leu SD medium containing 1 mM BPA. Cells were collected and resuspended in 1 ml of ice-cold PBS buffer. The cell suspension was then placed in a Spectrolinker XL-1500a (Spectronics Corp.) and irradiated by at 360 nm for 2×5 min. Extracts were prepared as described previously (Hu et al., 2011) and 5 mg of protein were incubated with 5 µl of Anti-PK antibody (Bio-Rad) for 2 hours at 4 °C. Next, 50 µl of Protein G Dynabeads (Life Technology) were added and incubated overnight at 4 °C to immunoprecipitate Scc1. After washing 5x with lysis buffer the beads were boiled in 2x SDS-PAGE buffer. Samples were run on a 3%–8% Tris-acetate gel (Life Technology) for 3.5 hours at 150 V. For Western blot analysis, Anti-Myc (Millipore) and Anti-FLAG (Sigma) antibodies were used to probe for Smc1 and Pds5, respectively.

### Coiled-coil predictions and conservation analysis

A set of SMC sequences and their domain delineations was used for coiled-coil prediction analysis (Bürmann et al., 2017). Individual coiled-coil probability profiles were generated with MARCOIL (Delorenzi and Speed, 2002), and both N- and C-terminal arm regions were extracted. N- and C-terminal profiles were separately aligned on their center coordinates, zero padded and averaged. We estimated 95 % confidence intervals for the averaged profiles using the 5 % and 95 % quantiles of 100 identically processed sequence sets generated by random resampling with replacement.

For conservation analysis, MukB sequences were aligned using MSAProbs (Liu et al., 2010). Jensen-Shannon divergences were computed for each alignment column according to (Capra and Singh, 2007), but using equal weights at positions with more than 30 % gaps.

## Data availability

Crystallographic structure factors and model coordinates will be deposited in the PDB.

## Code availability

The Xi software suite is available at https://github.com/Rappsilber-Laboratory/XiSearch. Custom code for statistical analysis is available on request.

## Author Contributions

Protein purification, F.B. and B.-G.L.; Electron microscopy, F.B. and B.-G.L.; Mass spectrometry and identification of cross-links, L.S., F.O.; CLMS data analysis and bioinformatics, F.B.; Crystallography, F.B. and J.L.; *E. coli* strain construction, F.B.; Yeast strain construction and *in vivo* cross-linking experiments, T.T.; Conception of the paper model, S.Y.; Preparation of the manuscript with input from all authors, F.B.; Supervision of the work, J.R., B.H., K.N., J.L.

## Acknowledgements

We are grateful to Danguole Kureisaite-Ciziene for help with crystallography and Minmin Yu for help with X-ray data collection. We thank Xian Deng, Francesca Coscia and Giuseppe Cannone for help with electron microscopy. We thank Julius Fredens for advice on recombineering and gift of the *pheS*\*-*hygR* cassette. We thank Gemma Fisher and David Sherratt for help with initial complementation experiments and gift of the *neoR* marker. F.B. is funded by an EMBO Long-Term Fellowship (EMBO ALTF 1151-2017). This work was funded by the Medical Research Council (U105184326 to J.L.), the Wellcome Trust (202754/Z/16/Z to J.L.), the DFG (25065445 to J.R.), and the Wellcome Trust through a Senior Research Fellowship to J.R. (103139). The Wellcome Centre for Cell Biology is supported by core funding from the Wellcome Trust (203149).

---

**Supplementary Figure 1.**
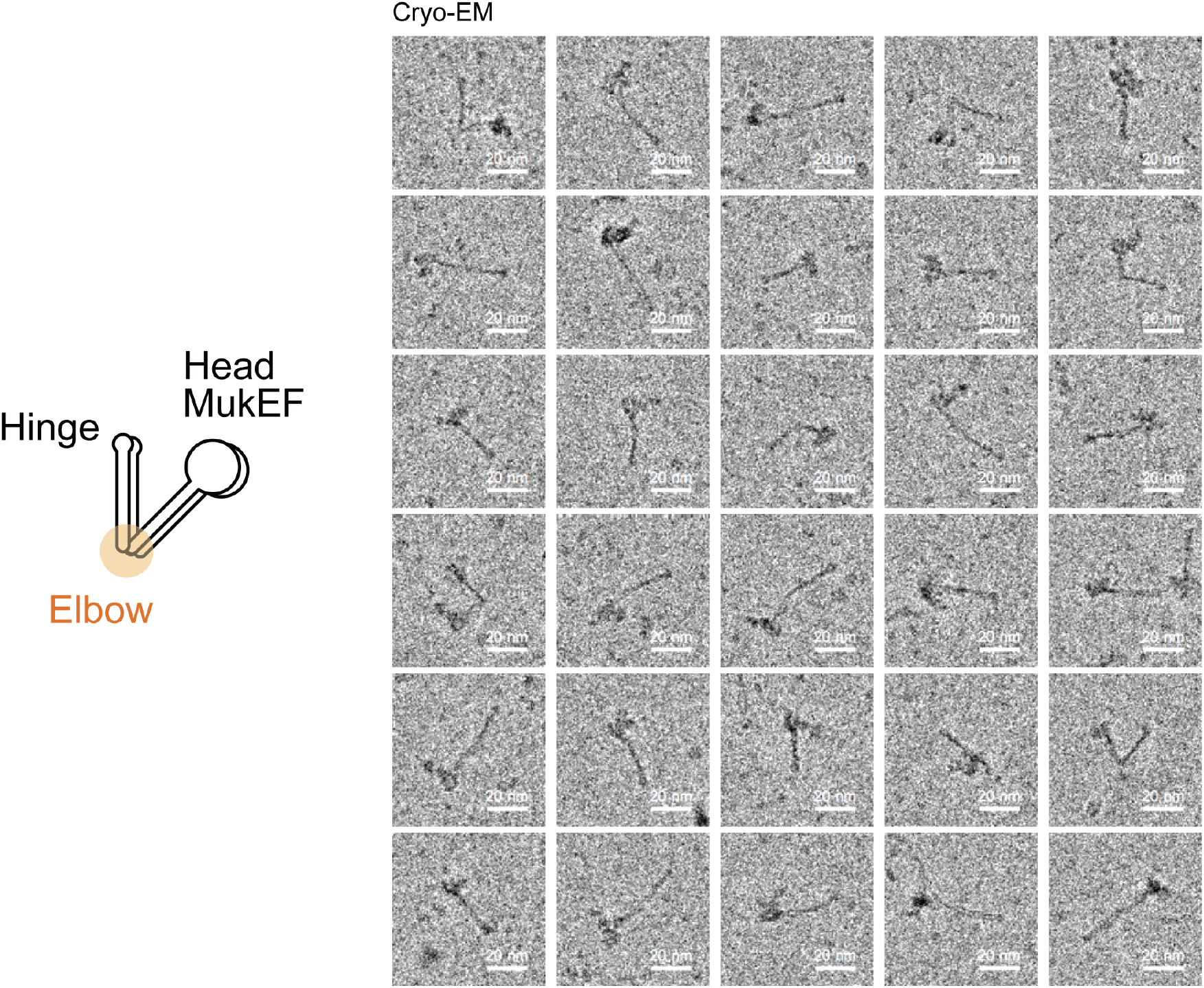
Cryo-EM of *Desulfovermiculus halophilus* MukBEF. Particles were imaged in unsupported vitreous ice and contrast was enhanced by use of a Volta phase plate. The presence of an elbow is indicated by a sharp central kink in the arm of several particles. Sequence identity between *D. halophilus* and *E. coli* MukBEF complexes is ~26 %.

**Supplementary Figure 2.**
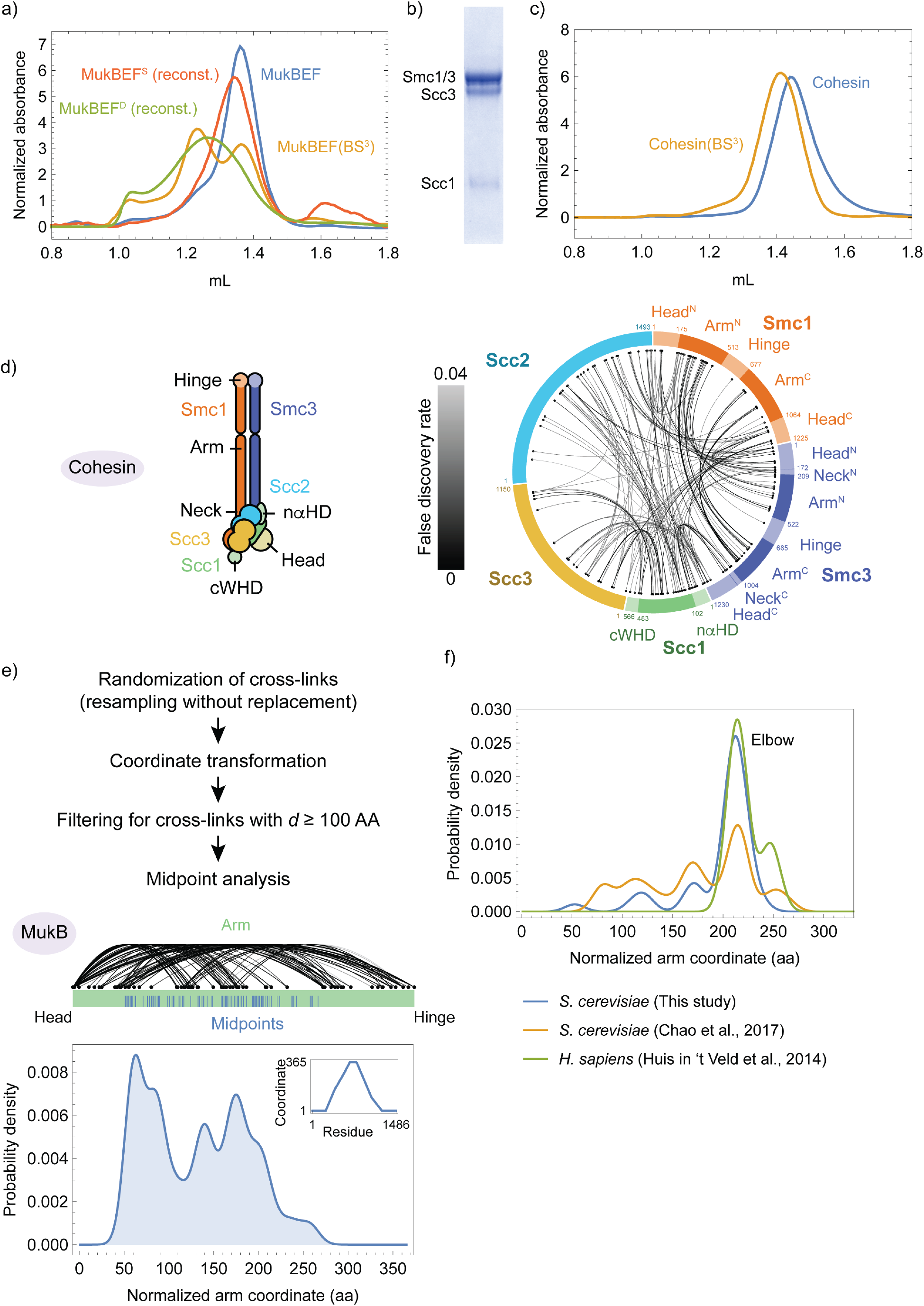
Cross-linking and mass spectrometry of MukBEF and cohesin. **(a)** SEC profiles of native co-expressed MukBEF (blue), BS3 treated co-expressed MukBEF (orange), singlet MukBEF (MukBEF^S^) reconstituted in buffer containing 40 mM NaCl, 2 mM MgCl2 (red) and doublet MukBEF (MukBEF^D^) reconstituted in buffer containing 200 mM NaCl (green). Reconstitution was similar to the protocol established in (Petrushenko et al., 2006). (**b**) SDS-PAGE analysis of a purified cohesin complex containing Smc1, Smc3, Scc1 and Scc3. The gel was stained with Coomassie. **(c)** SEC profiles of the cohesin complex containing Smc1, Smc3, Scc1 and Scc3 before and after treatment with BS3 (see Fig. 1g). **(d)** Inter-subunit cross-links of a cohesin complex containing Smc1, Smc3, Scc1, Scc3 and Scc2. As in Fig. 2a. **(e)** Cross-link midpoint analysis for MukB performed as in Fig. 2d but using random resampling without replacement before data processing. **(f)** Cross-link midpoint analysis for various cohesin datasets (as in Fig. 2). Peak density for human cohesin corresponds to residues 375/813 (Smc1) and 379/811 (Smc3), respectively.

**Supplementary Figure 3.**
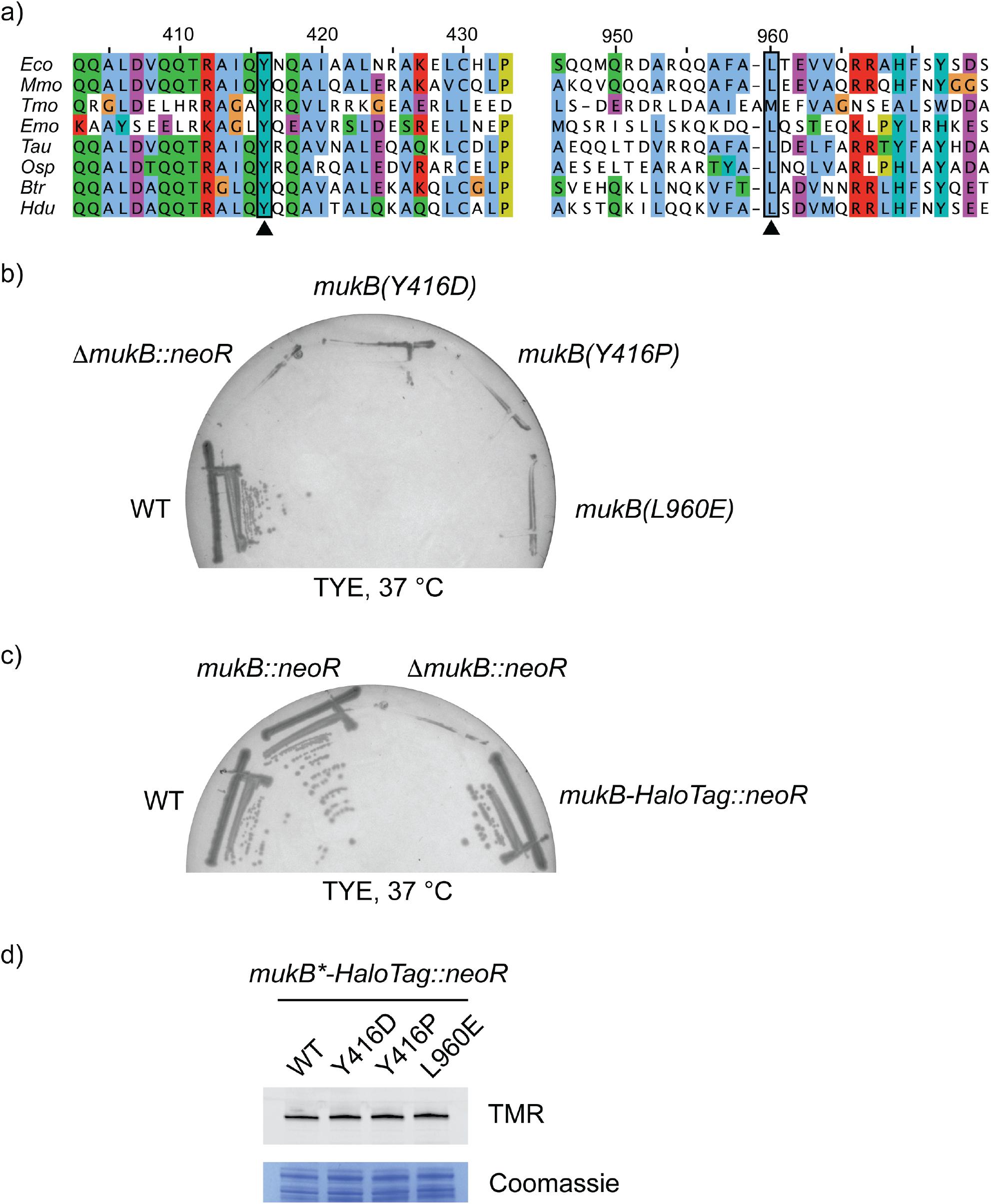
Mutagenesis of the MukB elbow. (**a**) Sequence alignment of the N-terminal (left) and C-terminal (right) parts of the MukB elbow. The mutated residues are highlighted by triangles. *Eco, Escherichia coli*; *Mmo, Morganella morganii*; *Tmo*, *Thioflavicoccus mobilis*; *Emo*, *Endozoicomonas montiporae*; *Tau*, *Tolumonas auensis*; *Osp*, *Oceanimonas sp*. GK1; *Btr*, *Bibersteinia trehalosi*; *Hdu*, *Haemophilus ducreyi*. (**b**) Growth of strains containing point mutations at the elbow in the endogenous *mukB* gene. (**c**) Construction of a functional *mukB-HaloTag* allele. (**d**) Protein levels of elbow mutants fused to a HaloTag. Extracts were labelled with a HaloTag-TMR substrate and were analyzed by in-gel fluorescence (top) and Coomassie staining (bottom) after SDS-PAGE. WT, wild-type.

**Supplementary Figure 4.**
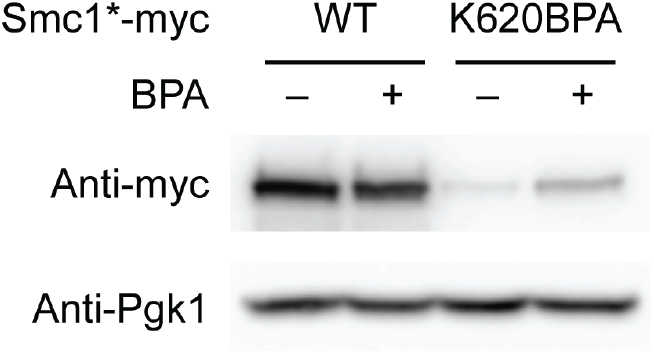
BPA-dependent expression of Smc1(K620BPA). Strains were grown either in the absence or presence of 1 mM BPA, and extracts were analyzed by Western blotting.

**Supplementary Figure 5.**
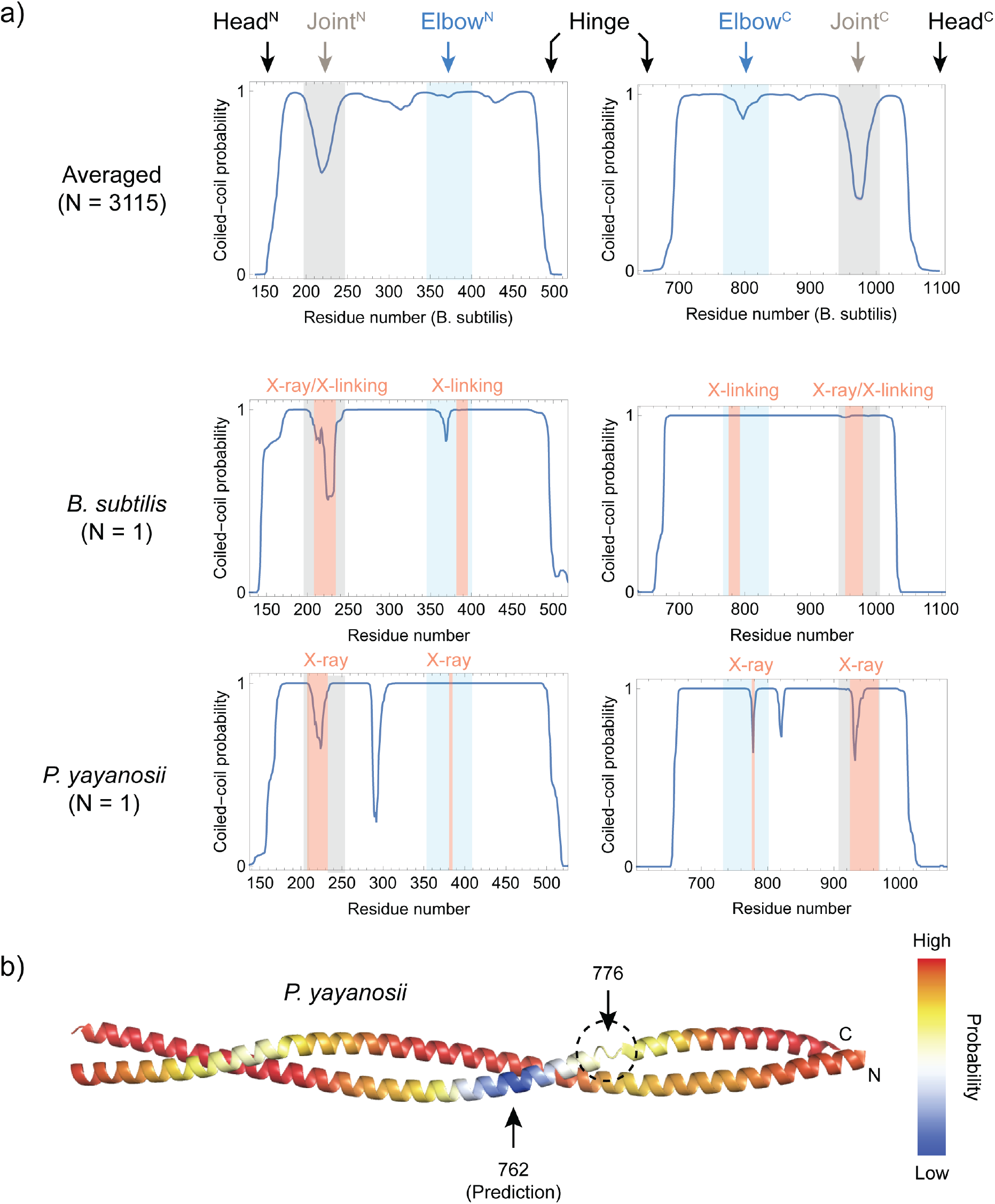
Locations of coiled-coil discontinuities in bacterial/archaeal Smc proteins. (**a**) Aggregate coiled-coil probability profile (same as in Fig. 5) and single-sequence profiles for *B. subtilis* Smc (bacterial) and *P. yayanosii* Smc (archaeal), respectively. Positions of coiled-coil discontinuities experimentally determined by X-ray crystallography (Diebold-Durand et al., 2017) or disulfide cross-linking (Waldman et al., 2015) are highlighted in red. (**b**) The elbow region of *P. yayanosii* Smc. The predicted coiled-coil probability from aggregate analysis (see **a** and Fig. 5) is mapped onto the crystal structure of a central arm fragment (PDB: 5XG2). Positions of the predicted and chrystallographically determined discontinuities are shown.

**Supplementary Figure 6.**
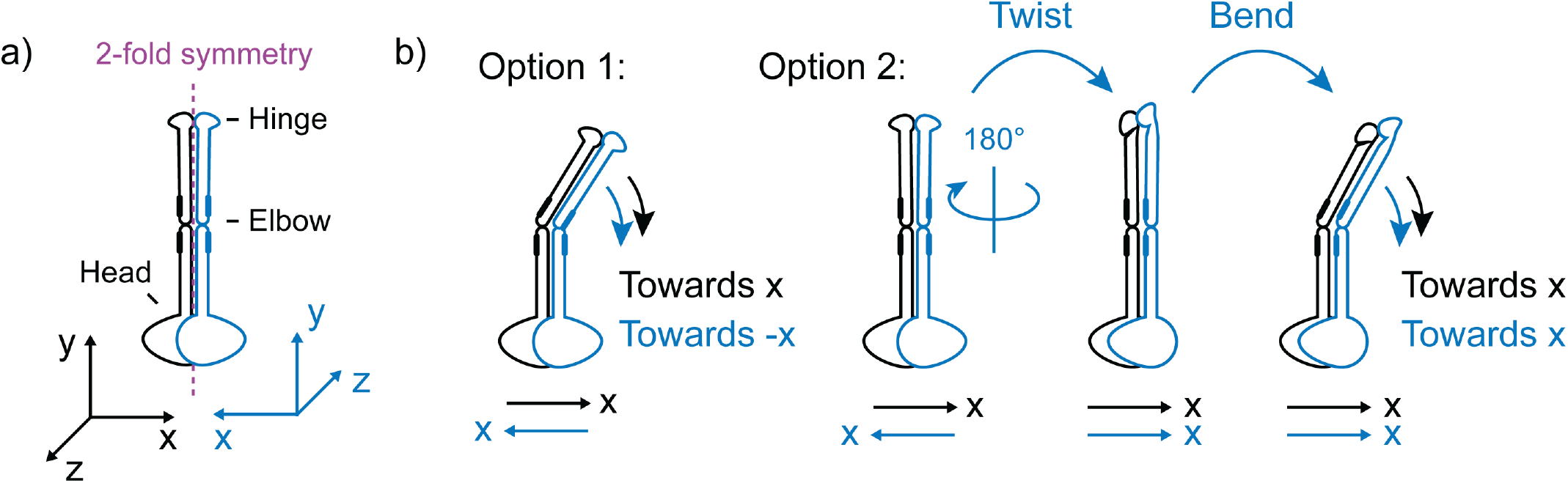
Bending of SMC dimers. **(a)** An SMC dimer with C2 symmetry. Monomers and their body-frame coordinate systems are shown in black or blue. The symmetry axis of the dimer is shown in purple. **(b)** Symmetry breaking upon elbow bending. Option 1: monomers bend into opposite directions; Option 2: monomers twist and bend into the same direction. Orientations of the relevant body-frame coordinate axes are shown at the bottom.

**Supplementary Figure 7.**
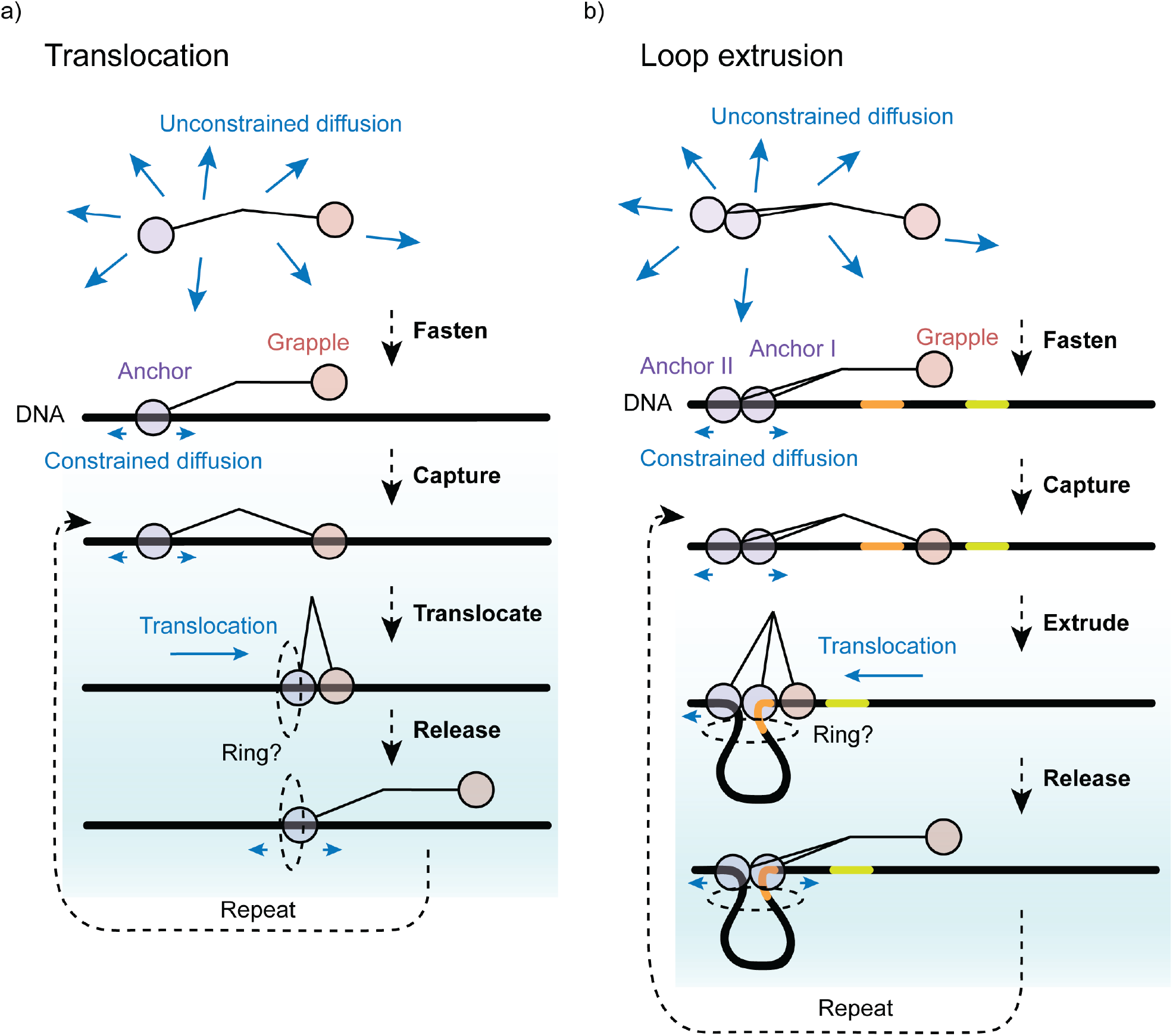
Detailed models for DNA and translocation and loop extrusion. (**a**) DNA translocation model requiring a regulated grapple DNA binding site and a sliding anchor DNA binding site. DNA binding may or may not involve a DNA entrapping ring that could be used to enhance processivity. **(b)** Loop extrusion using a second anchor site. DNA binding may or may not involve a DNA entrapping ring that could be used to enhance processivity.

## Supplementary Tables

**Supplementary Table 1** Cross-linking/mass spectrometry are available as a separate table online.

**Supplementary Table 2.**
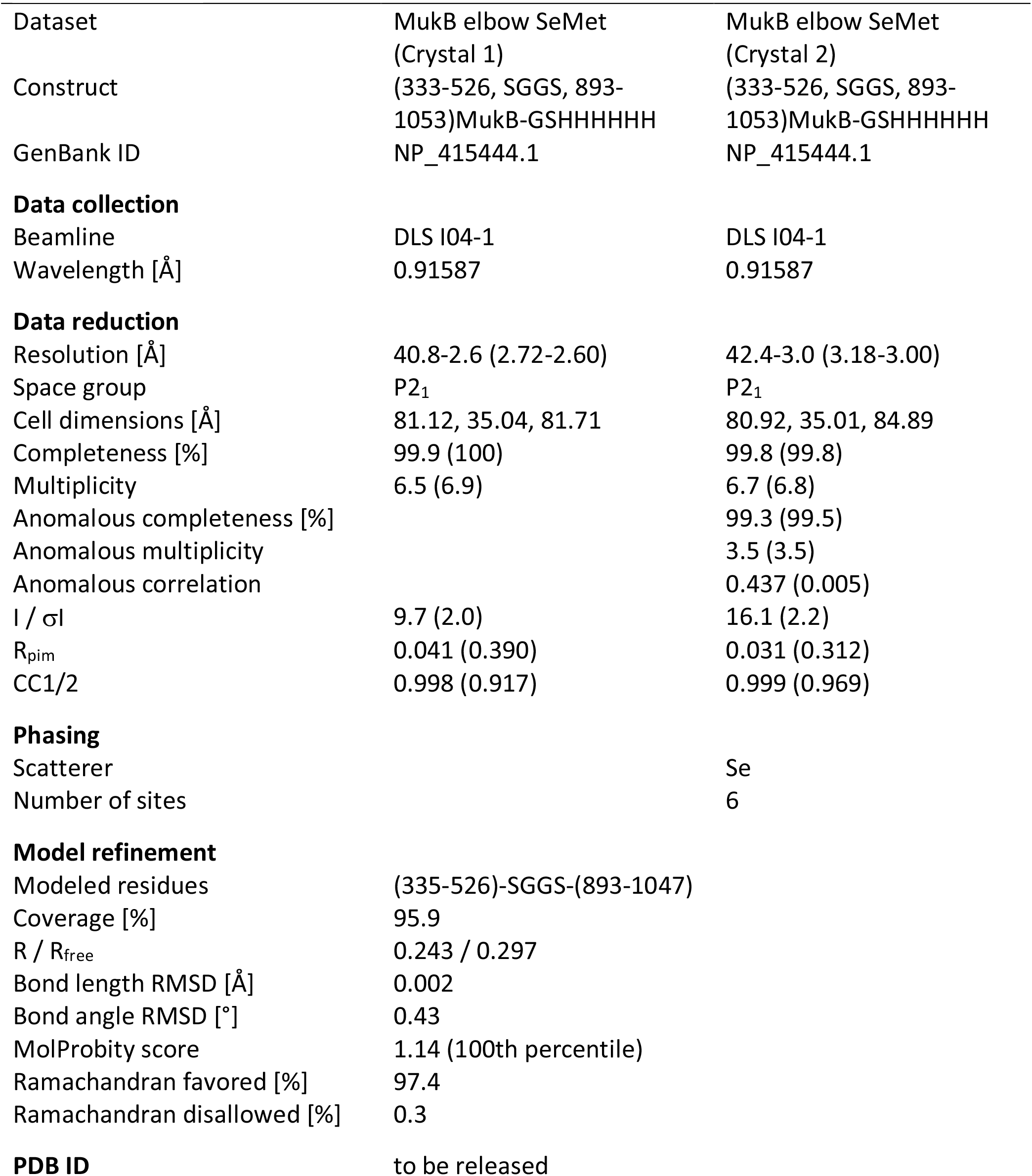
Crystallography table.

**Supplementary Table 3.**
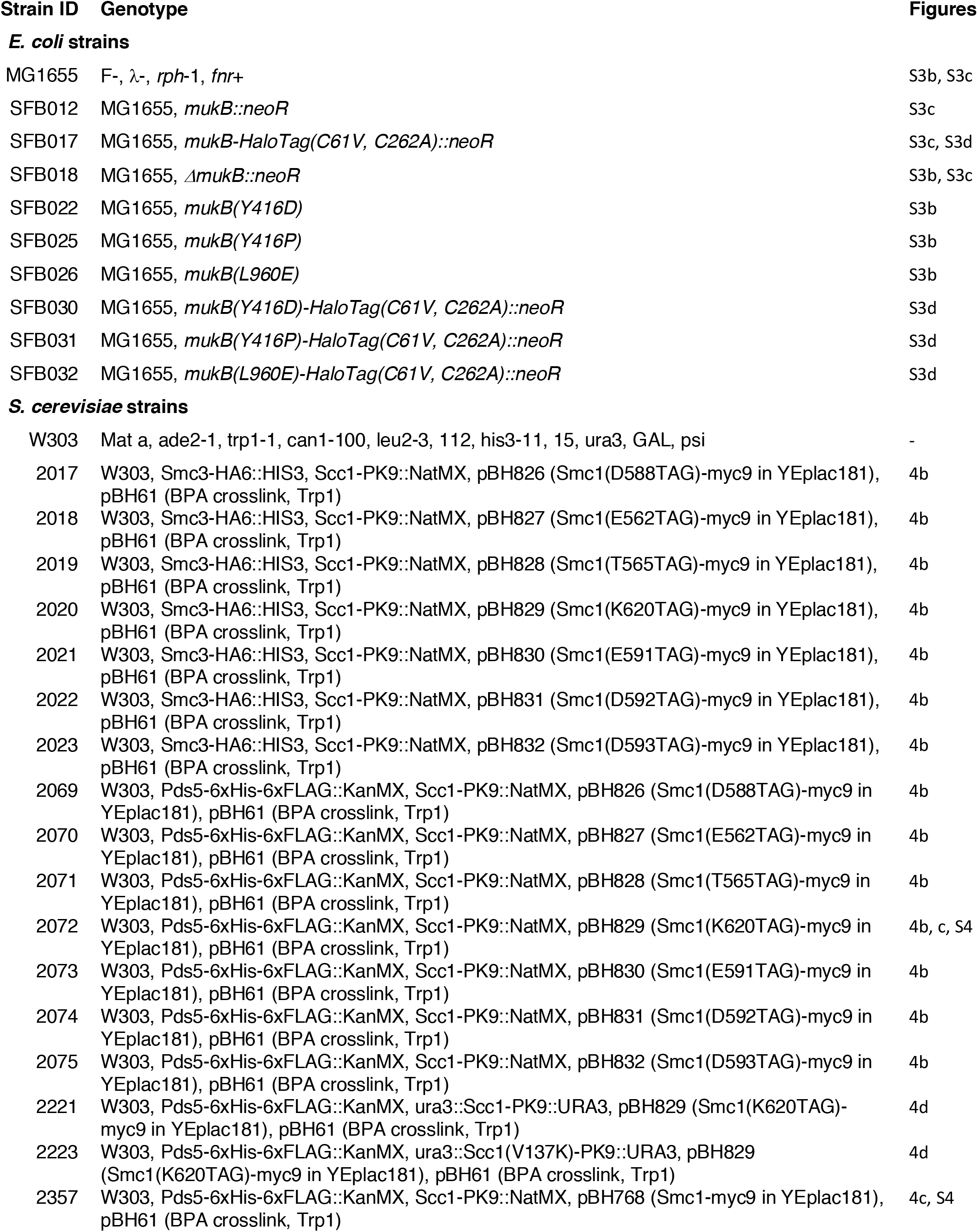

